# Model predicts the impact of root system architecture on soil water infiltration

**DOI:** 10.1101/2021.07.26.453789

**Authors:** Andrew Mair, Lionel X Dupuy, Mariya Ptashnyk

## Abstract

There is strong experimental evidence that root systems substantially change the saturated hydraulic conductivity of soil. However, the mechanisms by which roots affect soil hydraulic properties remain largely unknown. In this work, we made the hypothesis that preferential soil moisture transport occurs along the axes of roots, and that this is what changes a soil’s saturated hydraulic conductivity. We modified Richards’ equation to incorporate the preferential flow of soil moisture along the axes of roots. Using the finite element method and Bayesian optimisation, we developed a pipeline to calibrate our model with respect to a given root system. When applied to simulated root systems, the pipeline successfully predicted the pore-water pressure profiles corresponding to saturated hydraulic conductivity values, observed by Leung et al. (2018), for soils vegetated with willow and grass. Prediction accuracy improved for root systems with more realistic architectures, therefore suggesting that changes in saturated hydraulic conductivity are a result of roots enabling preferential soil moisture transport along their axes. The model proposed in this work improves our ability to predict moisture transport through vegetated soil and could help optimise irrigation, forecast flood events and plan landslide prevention strategies.

## 1 Introduction

Vegetated soils often exhibit higher saturated hydraulic conductivity values than fallow soils. Experimental evidence can be found in many scientific studies including the works of Archer et al. (2002), Wilcox et al. (2003), Scholl et al. (2014), Song et al. (2017) and Leung et al. (2018). However, numerous studies have also reported that the influence of root systems on the saturated hydraulic conductivity of a soil varies between different plant species and soil types (Gao et al., 2019). Song et al. (2017) observed that the saturated hydraulic conductivity of soil was significantly higher when vegetated with Vetivier grass rather than Bermuda grass. In fact, in some instances, they reported that Bermuda grass reduced moisture infiltration through the soil. These results indicate that not only root abundance but also the structure of the root system have an impact on the saturated hydraulic conductivity of the soil. However, the mechanism by which the soil properties are changed by the roots is not well characterised.

The root tissue itself tends to have lower hydraulic conductivity than soil. The low radial hydraulic conductivity of root tissues is a necessity for plants to resist drought and prevent the entry of toxic compounds or harmful pathogens. This low radial hydraulic conductivity is due to the composition of the cell wall, in particular the presence of Casparian strips which surround the walls of exterior root cells and mitigate the transfer of water and solutes into the tissue (Naseer et al., 2012). In contrast, root tissue is generally more conductive in the axial direction, in order to facilitate the transport of water and minerals to the shoots. These hydraulic properties were observed for maize root tissue in the experimental results of Frensch and Steudle (1989). Furthermore, the modelling work of Doussan et al. (1998) estimated that, at a distance of 60 cm from the tip of a primary maize root, the radial hydraulic conductivity would be in the range (0, 1] and the axial hydraulic conductivity would be in the range [4, 5], where the units of both values are m^4^s^-1^MPa^-1^ × 10^9^.

Based on these findings, it is therefore unlikely that the root tissue itself significantly enhances the saturated hydraulic conductivity of soil. It is widely accepted however, that the development of channels within a soil, through which moisture can preferentially flow, plays a significant role in altering soil’s saturated hydraulic conductivity (Scholl et al., 2014; Sidle et al., 2001). The decay and shrinkage of older roots in an architecture is known to open up pathways in soil which cause preferential transport of moisture in directions parallel to old or dead roots (Gile et al., 1995). Furthermore, it was reported by Dorioz et al. (1993) that in the rhizosphere, adjacent to living roots, the proximity of clay particles to one another increased as a result of roots penetrating through the soil. The compactness of the rhizosphere is further increased by the chemical action of the mucilage that is exuded by growing roots (Carminati et al., 2010; Johnson and Lehmann, 2006). This opens up space in the nearby non-rhizosphere soil and makes it more porous (Angers and Caron, 1998). The observations of Noguchi et al. (1997) support these findings by showing that living roots blocked local moisture flow and redirected it around their perimeters into neighbouring regions of more conductive soil.

It is likely, therefore, that pathways in a soil will develop parallel to the axes of both living and dead roots, and that a change in the soil’s saturated hydraulic conductivity is a result of preferential flow through these pathways. Unfortunately, experimental studies of preferential soil moisture transport along root axes are lacking (Ghestem et al., 2011). The saturated hydraulic conductivity of a vegetated soil is traditionally measured by first applying a constant ponding head to a column of saturated vegetated soil, then calculating the saturated hydraulic conductivity as the steady state infiltration rate divided by the hydraulic gradient (Leung et al., 2018). This approach only considers vertical moisture transport, and the contribution of preferential flow along individual roots is neglected. The more modern approach of Electrical Resistivity Tomography does allow three-dimensional moisture distributions in vegetated soils to be obtained, and these provide richer information about the impact of root structure and preferential flow on the hydraulic properties of vegetated soil (Beff et al., 2013). Such techniques require specialised equipment however and, at present, access to these facilities is limited.

There were two main objectives of this study. The first objective was to develop a new model for moisture transport through vegetated soil which incorporated preferential soil moisture flow along root axes. The second objective was to use the model to test the hypothesis that root-induced preferential flow of soil moisture is the main cause of the difference in saturated hydraulic conductivity between vegetated and fallow soils. We integrated our model into a software pipeline which linked root system simulation, volumetric root density construction, finite-element approximation of partial differential equations, and model calibration to predict three-dimensional soil moisture and pore-water pressure profiles within vegetated soils. Using this pipeline with the experimental data of Leung et al. (2018), we provided evidence that preferential soil moisture transport along root axes is a key cause of the differences in saturated hydraulic conductivity between varyingly vegetated soils.

## 2 Mathematical models for root-induced preferential flow

Richards’ equation, (Richards, 1931),

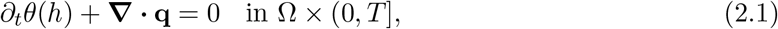

is the well established continuity equation for moisture transport through fallow unsaturated soil. Here 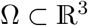 is a Lipschitz domain representing a volume of soil and *T* > 0 is a final time considered in experiments or numerical simulations. The flux q in (2.1) is proportional to the negative gradient in hydraulic potential and is given by Darcy’s law, (Darcy, 1856),

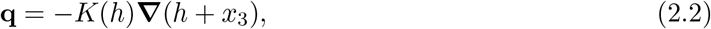

where *x* = (*x*_1_, *x*_2_, *x*_3_) ∈ Ω and *x*_3_ is pointing in the positive vertical direction. The variable *h* is the pressure head (m) and is related to pore-water pressure *p* (Pa) by the formula *p* = *ρgh*, where *ρ* is the soil moisture density (kgm^-3^) and *g* is gravitational acceleration (ms^-2^. The two constitutive relations in (2.1) and (2.2) are the soil moisture retention function for soil moisture content *θ* (—) and the soil hydraulic conductivity *K* (ms^-1^). The Van Genuchten (1980) formulas

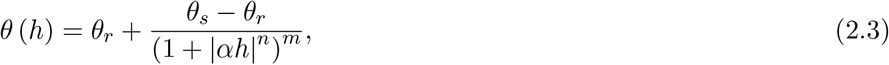

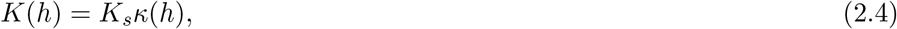

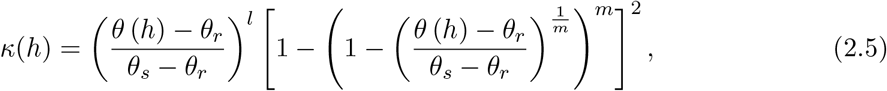

are the most popular choices in experimental studies of groundwater flow and theoretical analysis of Richards’ equation, see e.g. Liang et al. (2017), Leung et al. (2018), Radu et al. (2008) and Radu et al. (2018). The parameters *θ_r_* and *θ_s_* in (2.3) are the residual and saturated moisture contents (—) respectively, and *α* (m^-1^), *n* (—) and *m* = 1 —1/*n* are all shape parameters that take different values depending on the soil type under consideration. In (2.4), the parameter *K_s_* is the saturated hydraulic conductivity (ms^-1^) of the soil being considered. The function *κ* in (2.5) defines the constitutive relation between hydraulic conductivity and pressure head *h*, where the constant *l* is the pore connectivity (—) of the soil. For ease of notation these parameters have all been stated with time in seconds (s), however, for the models, methods and results of this paper, time shall hereafter be given in days (d).

To incorporate the influence of a root system 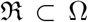 on soil moisture transport, we assume that the root system can be expressed as a union of 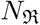 conical frustums 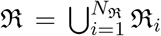 and modify the flux of Richards’ equation (2.1) to develop our ‘facilitation model’. In this model the influence of 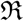 on the transport of soil moisture through Ω depends upon the length, radii and orientation of each root segment in 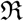. To introduce the model we first consider the case of a homogeneous root system 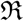 where all segments have the same dimensions and orientation. Assuming that there is a maximal lateral surface area *S*_max_, for a single root segment from any root system, we define the single normalized lateral surface area exhibited by each segment in our homogeneous root system as 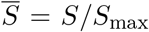, where *S* is the lateral surface area common to each segment in this system. By the design of our model, the root system 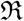 induces anisotropy in the hydraulic conductivity of the soil. As a result, the following matrix is used to distinguish between the radial and axial influence of the segments of the root system

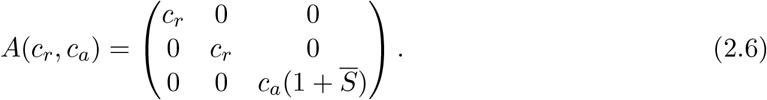

Here *c_r_* > 0 is the radial facilitation coefficient and *c_a_* > 0 is the axial facilitation coefficient. If *c_a_* > 1 then the macroscopic hydraulic conductivity is increased in the axial direction of roots, and if *c_a_* < 1 then it is decreased, the same applies to the radial direction of roots regarding the value of *c_r_*. In this work we focused on the hypothesis that preferential flow occurs axially which meant *c_r_* = 1 and *c_a_* > 1. To account for the orientation of each segment in the homogeneous root system 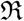, the matrix in (2.6) undergoes the following transformation

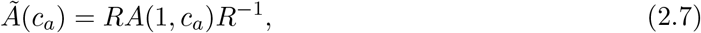

where the operation *R*(0,0,1)^⊤^ produces the unit vector parallel to the direction of root segments in 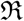. The rotation matrix *R* is computed via the method of Rodrigues (1840) and more details are in appendix A. The preferential moisture transport along the axes of segments in 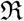 is then incorporated into (2.1) by replacing q in (2.2) with

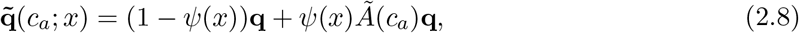

where *ψ* is the volumetric root density function for the system, providing a continuous approximation of the position, size and orientation of the segments. The first term on the right hand side of (2.8) represents the flux of moisture through bulk soil where root segments have no influence, and the matrix 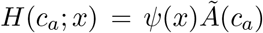 in the second term incorporates preferential transport of soil moisture along the axes of the segments of 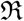.

In reality root systems are heterogeneous and comprised of segments of differing size and orientation. As a result, the volumetric root density *ψ* and heterogeneity matrix *H*(*c_a_*, *x*) for an heterogeneous root system are formulated by summing the influence of each individual segment 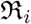 in the root system 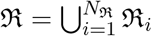. For a segment 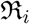 the centres and radii of the circular ends are *a_i_*, *b_i_*, and *r*_a_i__, *r*_b_i__ respectively. In addition the length, midpoint, volume and lateral surface area of 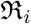 are denoted by 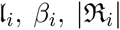 and *S_i_* respectively, with the normalised lateral surface area being given by 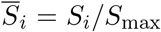. The volumetric density *ψ_i_* which corresponds to 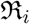 is then defined as

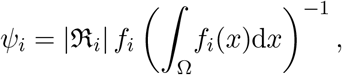

where *f_i_* is the probability density function of the multivariate normal distribution 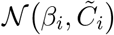. The covariance matrix of this distribution is defined as 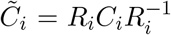 where *R_i_*(0, 0,1)^⊤^ produces the unit vector parallel to 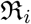 and

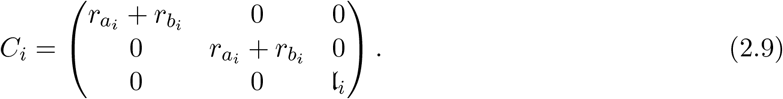

This construction means that *ψ_i_* provides a continuous approximation of the position, size and orientation of the segment 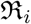 with 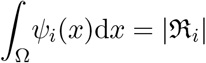. Given an axial facilitation constant *c_a_* for the entire system 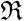, the influence of the segment 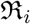, on the flux of moisture transport in Ω, is incorporated using its associated heterogeneity matrix 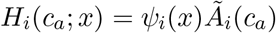 for *x* ∈ Ω, where for each segment 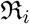 we have 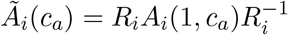 and

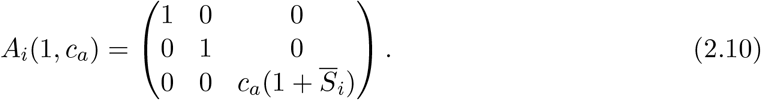

The volumetric density function *ψ*: Ω → [0,1] and heterogeneity matrix 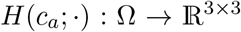, for the entire heterogeneous root system, are then defined as 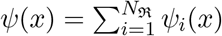 and 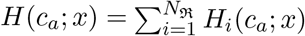 for *x* ∈ Ω. By this construction *ψ* gives a continuous approximation of the entire root system and the matrix *H* incorporates the preferential flow along the axis of every segment in the root system. The moisture flux term which incorporates preferential flow along the axes of all roots in the system is then given by

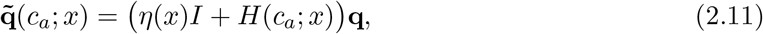

where *η*(*x*) = 1 — *ψ*(*x*) is the complement of the volumetric root density. This means that in regions of high root abundance, preferential flow is induced along the axes of all nearby root segments and the soil hydraulic conductivity in (2.11) is materially influenced by the root system. In regions of low root abundance the hydraulic conductivity in (2.11) is essentially the isotropic Darcy flux (2.2) and the root system has little influence on soil moisture transport. The full ‘facilitation’ equation which incorporates, into (2.1), the preferential flow of soil moisture along the axes of a root system 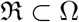 is given by

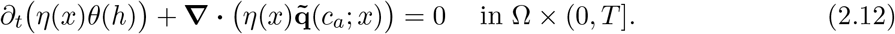

Despite the hypothesis that root segments facilitate the flow of soil moisture along their respective axes, the fact remains that root tissue itself is an obstacle to soil moisture transport. This notion is incorporated into (2.12) through the additional factor of *η*(*x*) in the time derivative and flux. In numerical simulations and analysis of (2.12) we considered a soil column domain 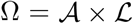, where 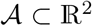 has Lipschitz continuous boundary *∂A* and 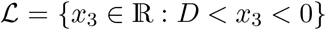, with *D* being the depth of the soil column.

To calibrate (2.12) for a given root system 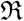 we used the experimental data, see Table 1, and the ‘empirical’ model of Leung et al. (2018) given by

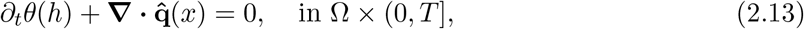

with

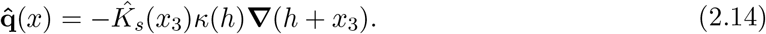

**Table 1:** Rooting depth of each age class and species of plant along with the corresponding effective saturated hydraulic conductivity 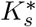 of the soil that they occupied. These values were obtained experimentally by Leung et al. (2018).

| Species | Willow |  |  |  | Grass |  |  |  |
| --- | --- | --- | --- | --- | --- | --- | --- | --- |
| Plant age (weeks) | 2 | 4 | 6 | 8 | 2 | 4 | 6 | 8 |
| Rooting depth, $\vert x_3^*\vert $ (m) | 0.256 | 0.450 | 0.630 | 0.800 | 0.172 | 0.367 | 0.450 | 0.500 |
| $K_s^*$ (md<sup>-1</sup>) | 0.435 | 0.860 | 1.503 | 2.212 | 0.187 | 0.530 | 0.660 | 1.140 |

Here the influence of a root system 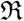 on the moisture transport through a soil column Ω is incorporated by considering in Richards’ equation (2.1) the depth dependent saturated hydraulic conductivity

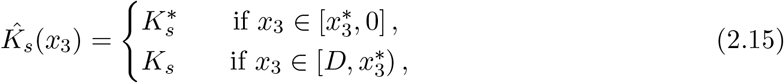

instead of the constant saturated hydraulic conductivity *K_s_*. In (2.15) the value 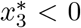 is the depth that the root system 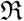 reaches within the soil column and the constant 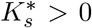 is the experimentally obtained value for the effective saturated hydraulic conductivity of the vegetated soil that lies above the depth 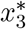. The specific values for 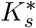 which Leung et al. (2018) recorded for soil columns vegetated with willow and Festulolium grass plants of different ages are shown in Table 1.

**Figure 1:**
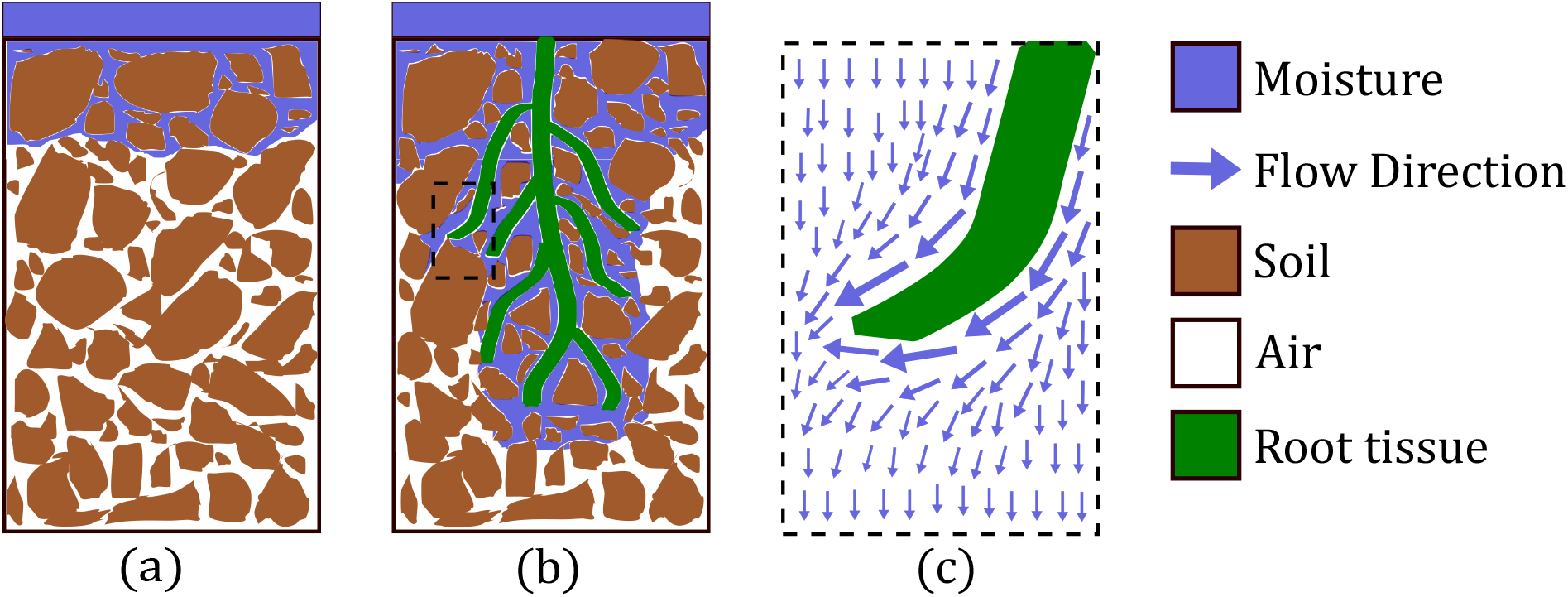
An illustration of the hypothesis that vegetation increases the hydraulic conductivity of soil, specifically by inducing preferential moisture transport along the axes of roots.

To formulate the full ‘facilitation’ and ‘empirical’ models, we equip (2.12) and (2.13) with the following boundary and initial conditions

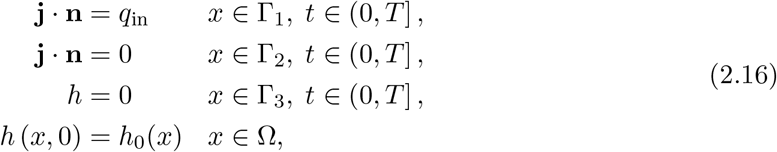

where **j** denotes the flux of the equation being considered and **n** denotes the outward unit normal vector, i.e. 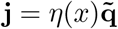 for (2.12) and 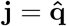 for (2.13). The soil-atmosphere boundary of the domain is denoted by 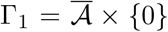, where 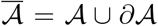. The lateral boundary is 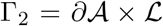 and the lower soil-soil boundary is 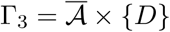. The Neumann boundary condition *q_in_* < 0 on Γ_1_ reflects the rate of moisture infiltration into the soil and we assume zero flux conditions on the lateral surfaces of Ω. When equipping (2.13) with (2.16), the infiltration condition is *q_in_* = *g_in_* but for (2.12) we have *q_in_*(*x*) = *η*(*x*)*g_in_* for *x* on Γ_1_. The Dirichlet condition on Γ_3_ reflects the assumption that Γ_3_ is located at the interface between the soil and the water table.

### 2.1 Model for simulating root system data

Equation (2.13) does not model the explicit influence of a root system’s architecture on soil hydraulics. However, since the saturated hydraulic conductivity values, 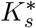, used in (2.13) result directly from experiments performed on soil vegetated by a root system, its solution profiles can be used as a benchmark to test and calibrate (2.12). This involves parametrising (2.12) by constructing the functions *ψ* and *H* which correspond to each of the willow and grass root systems examined by Leung et al. (2018). As demonstrated above, carrying out these constructions requires root system data in the form of a union of segments 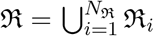, where the position, orientation and size of each segment are available. Such detailed data was not recorded by Leung et al. (2018), instead they measured the total root length, rooting depth and distribution of segment diameters for each of the willow and grass plants they investigated. We therefore use an algorithm to simulate root systems 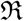 with physical properties that agree with these measurements. Our algorithm constructs a matrix 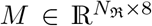, where for 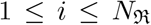 the entries (*M*_*i*,1_, *M*_*i*,2_, *M*_*i*,3_) ∈ Ω are the centre point of the proximal end of 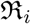, the entry *M*_*i*,4_ is the diameter of the proximal end, the entries (*M*_*i*,5_, *M*_*i*,6_, *M*_*i*,7_) ∈ Ω are the centre point of the distal end of 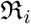 and *M*_*i*,8_ is the diameter of the distal end. The construction process for *M* is encapsulated by Algorithm 1. In the simulated root systems, for each segment 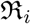 the azimuthal angle *ϑ_i_* is drawn from the uniform distribution *U*(0, 2π) whereas the polar angle 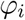 from the positive *x*_3_-axis is either drawn from the uniform distribution 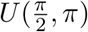 or from the truncated normal distribution 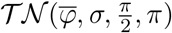. Here 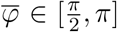 and 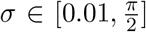 are the mean and standard deviation of the parent normal distribution 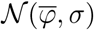 respectively, with 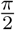 and π in 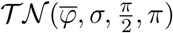 specifying the truncation interval (Burkardt, 2014). In the case where the polar angle is drawn from the truncated normal distribution 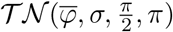 we have

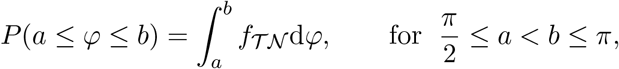

where

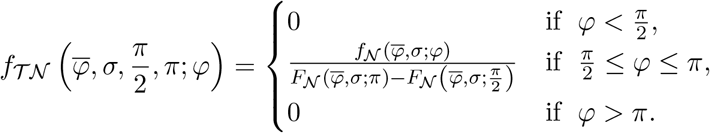

Here 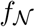 and 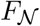 are the probability and cumulative density functions of the parent normal distribution 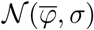 respectively. When a uniform distribution is used for 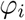, the resulting root system is referred to as uniform and when a truncated normal distribution is used, the root system is referred to as truncated normal with parameters 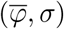. The cost functions and optimisation routine that are then used to calibrate (2.12), (2.16) in accordance with the data in Table 1 and the corresponding empirical solution profiles from (2.13), (2.16) are detailed in Section 3.

#### Algorithm 1: Root System Simulator

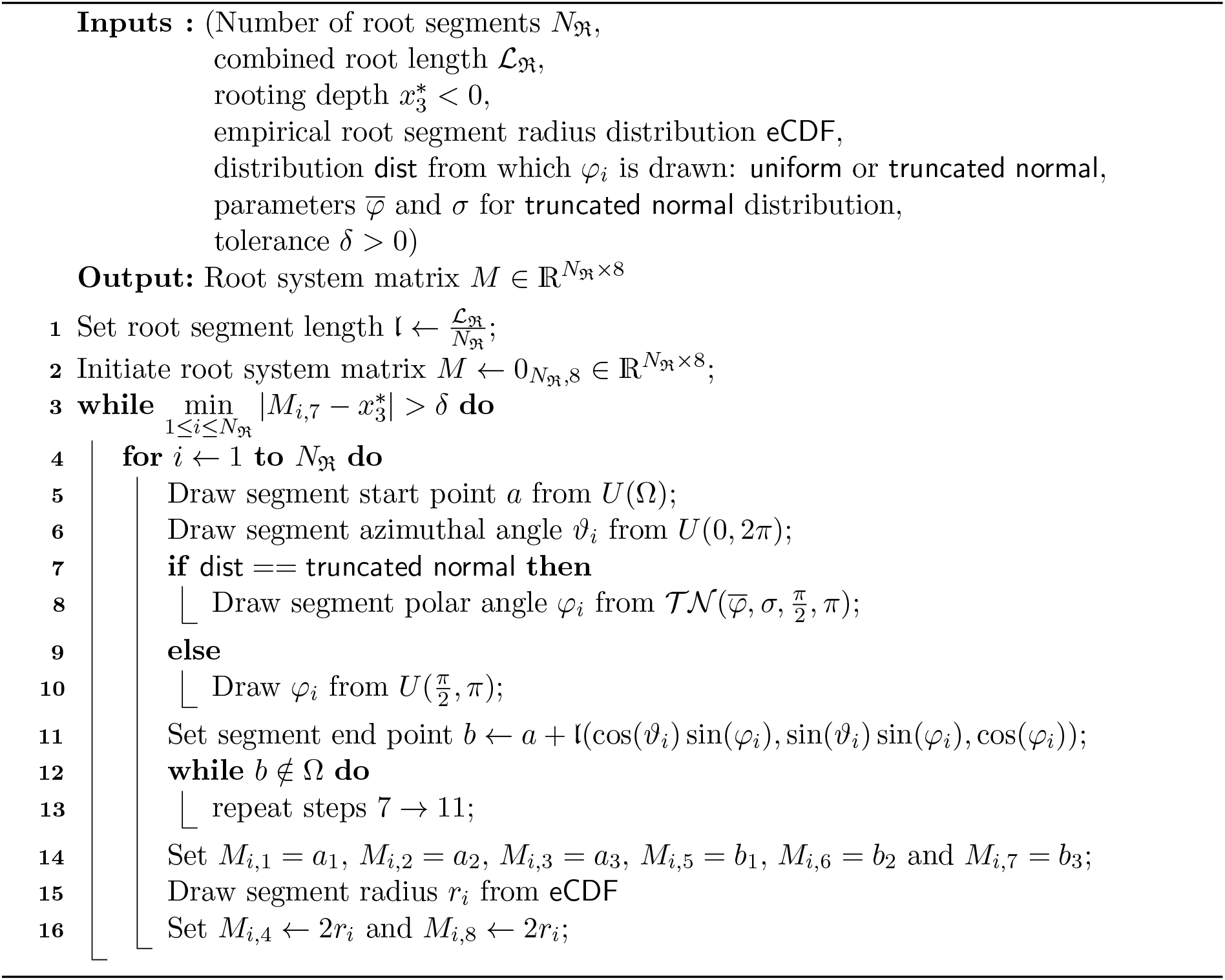

### 2.2 Numerical method for simulations of empirical and facilitation models

To obtain numerical solutions to models (2.13), (2.16) and (2.12), (2.16), we use the first-order conformal finite element method, together with the L-scheme for the linearisation of functions *θ* and *κ*, see e.g. (List and Radu, 2016; Slodicka, 2002). To discretise in time, 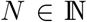 is set as the number of time steps and the step size is *τ* = *T/N*. Then for *n* ∈ 1,…, *N* with *t_n_* = *τn* and *h^n^* = *h*(·, *t_n_*), the time derivative is approximated using the backward Euler method

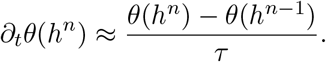

Applying the L-scheme, which exploits the Lipschitz continuity of *θ*, we write *θ*(*h^n^*) in the following form

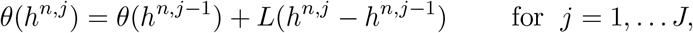

where *J* is the total number of iterations, *h*^*n*,0^ = *h*^*n*-1^, and L ≥ L_*θ*_ = sup_*h*≥_ |*θ*’(*h*)|, see List and Radu (2016). The time discretised and linearised variational form of (2.12), (2.16) then reads as follows: for *n* ∈ {1,…,N} and *j* ∈ {1,…*J*} find 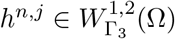 such that

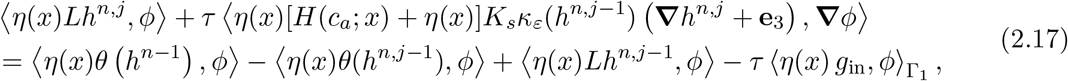

for all 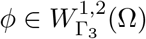. Here we use the notation 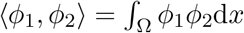 and 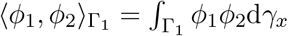, along with 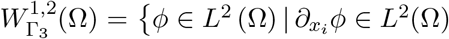, for *i* = 1, 2, 3, *φ* = 0 on Γ_3_}, where *L*^2^(Ω) is the space of all square integrable functions on Ω. The discretisation in space is achieved by projecting (2.17) onto a first-order continuous Galerkin finite element space *V_h_*. The space *V_h_* is defined by the decomposition of Ω into a finite family of disjoint simplexes 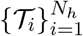 so that 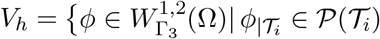, for *i* = 1,…,*N_h_*}, where 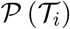 is the space of first-order polynomials on 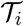, see e.g. Thomée (2006) for more details on continuous Galerkin finite element discretisations. This numerical scheme was implemented in Python 3 using the FEniCS library (Alnæs et al., 2015).

In (2.17) we use a regularisation *κ_ε_* of (2.5) to satisfy conditions on *θ* and *κ* which ensure the convergence of the numerical method used here, see e.g. (List and Radu, 2016). These conditions are as follows:

(A1) The retention function θ(·) is non-decreasing and Lipschitz continuous.
(A2) The function *κ*(·) is Lipschitz continuous and there exist *κ*_min_ and *κ*_max_ such that 0 < *κ*_min_ ≤ *κ*(*h*) ≤ *κ*_max_ < ∞, 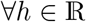.

For proofs that *θ*(·) from (2.3) satisfies (A1) see appendix C. The exact formulation of the regularisation *κ_ε_*(*h*) and verification of (A2) can be found in appendix D. In addition to these conditions, there exists *η*_0_ > 0 such that 0 < *η*_0_ ≤ *η*(*x*) ≤ 1 for *x* ∈ Ω. Furthermore, for any root system 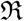 and axial facilitation constant *c_a_* ≥ 1, the heterogeneity matrix *H*(*c_a_*; *x*) in (2.12) is symmetric and uniformly positive definite, see appendix B for the proof.

## 3 Parametrisation and calibration

### 3.1 Incorporating experimental data into the facilitation model

Models (2.12), (2.16) and (2.13), (2.16) are parametrised with respect to the experimental measurements of Leung et al. (2018) for the saturated hydraulic conductivity of soils vegetated by willow and Festulolium grass (Table 1). The soil column domain 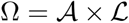 has a depth of 2m and a diameter of 0.025 m, i.e. 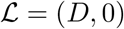 with *D* = —2 and 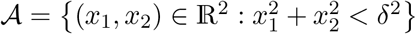 with *δ* = 0.025. The flux at the top boundary of the soil, modelling water infiltration, is *g*_in_ = —0.01 md^-1^, the saturated and residual water contents are *θ_s_* = 0.383 and *θ_r_* = 0.17 respectively, and the shape parameter values are *α* = 1.47 m^-1^ and *n* = 1.43. The saturated hydraulic conductivity for fallow soil in (2.13), (2.16) and (2.12), (2.16) is set to *K_s_* = 0.187 md^-1^, and the effective saturated hydraulic conductivity values 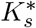 in (2.13), (2.16), observed experimentally, are given in Table 1. The initial pressure head profile in both models is set to 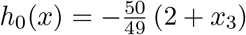.

The functions *ψ* and *H* are determined for a given root system by applying the methods detailed in Section 2.

Since root architecture data were not recorded by Leung et al. (2018), we used Algorithm 1 to simulate 8 week old willow and grass root systems which respected the measurements that they did record for each of these root systems. We initially use Algorithm 1 to simulate 8 week old willow and grass root systems where, for each segment, the angles on the horizontal plane and from the upward vertical axis are both drawn from uniform distributions. In the measurements of Leung et al. (2018) the 8 week old willow root system had a smaller total root length than the 8 week old grass, yet the willow root system reached greater depths within the soil column. To reflect this, we also simulate 8 week old willow and grass root systems where the root segments are oriented in a more downward direction in the willow root system and in a more horizontal direction in the grass root system. This is done by drawing the angle from the upward vertical axis of each segment from a truncated normal distribution where the parent normal distribution has mean 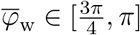 for the willow root system and mean 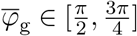 for the grass root system. See Section 2.1 for a detailed explanation of how grass and willow root systems are simulated. With the functions *ψ* and *H*, the facilitation model (2.12), (2.16) can be parametrised for each of the simulated root systems. Calibration of (2.12), (2.16) with respect to the experimental data in Table 1 then requires the minimisation of a cost function which expresses the difference between the solutions of the facilitation model (2.12), (2.16) and the solutions of the corresponding empirical model (2.13), (2.16). The cost function and variations thereof are detailed in Section 3.4 and the optimisation algorithm used for their minimisation is described in Section 3.2.

### 3.2 Optimisation

The cost functions, that we are required to minimise in order to calibrate the facilitation model (2.12), (2.16) are formulated in terms of numerical solutions to the facilitation and empirical models. Therefore, they cannot be expressed analytically, their derivatives cannot be obtained easily and we have no knowledge of their convexity properties. A classical approach to minimising functions such as these is Powell’s method which is a line search algorithm (Brent, 2013). Evaluations of the cost functions involve time intensive steps, such as the construction of heterogeneity matrices *H* and simulation of numerical solutions to nonlinear PDEs. As a result, Powell’s method often requires unattractively long computation times to find minimisers, especially since the number of cost function evaluations performed cannot be set a priori. This motivates the use of Bayesian optimisation which does not require derivatives of the cost function and efficiently explores the parameter space. Furthermore, the number of function evaluations that are performed by a Bayesian optimisation algorithm can be fixed a priori.

Bayesian optimisation is based on Bayes theorem (Williams, 1991) and uses two components to search for the minimiser of a cost function 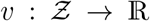, where 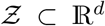 is the parameter space with dimension *d* > 0. The first component is the surrogate probability model 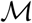 which is used as the prior distribution for the value of a cost function *υ*(**z**) evaluated at a parameter value 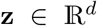. Given *n* observations of the value of the cost function, 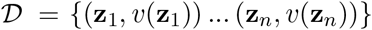, the surrogate 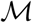 is conditioned on 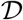 using Bayes theorem. The second component of Bayesian optimisation is the acquisition function 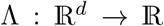, which is defined by the choice of surrogate 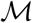 and depends upon the current contents of the observation set 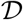. This function Λ proposes the best location 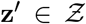 at which to evaluate *υ*, so as to most efficiently find its minimiser **z**^*^. The proposed next evaluation location for any choice of acquisition function Λ is the value **z**’ which maximises Λ(**z**’) (Brochu et al., 2010). The full Bayesian optimisation routine of N**f**_evs_ function calls is given by Algorithm 2.

#### Algorithm 2: Bayesian optimisation (Dewancker et al., 2016)

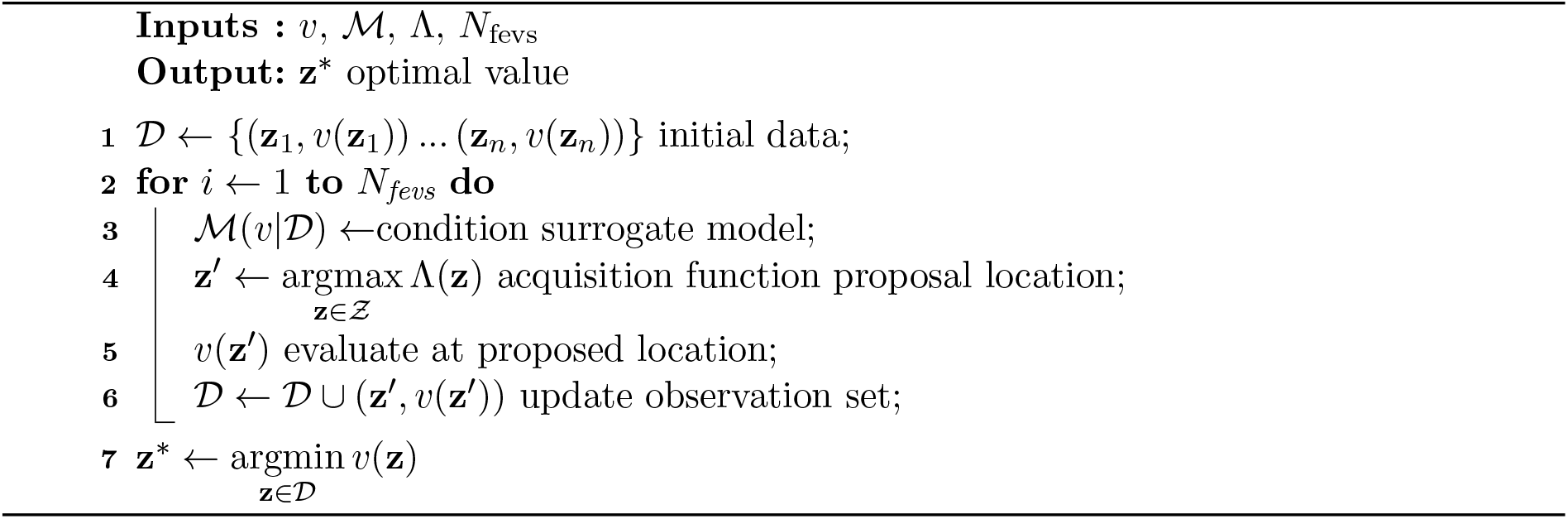

There are a number of options for the choice of surrogate model 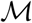 and acquisition function Λ, see e.g. Brochu et al. (2010), but in this work we opted for the combination of a Gaussian process 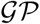 surrogate model and an expected improvement acquisition function:

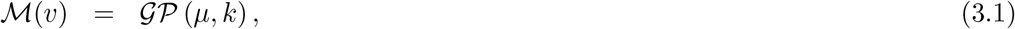

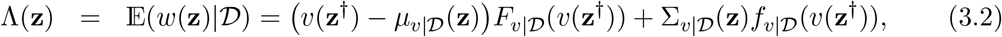

where 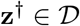 is the best minimiser of *υ* that has been observed so far and

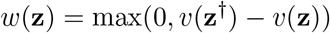

is the so-called improvement function. The functions 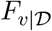 and 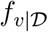 are the cumulative and probability density functions of the conditioned distribution 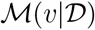 respectively. By the definition of a Gaussian process, any finite vector of *n* observations **v**_n_ = (*υ*(**z**_1_),…,*υ*(**z**_n_))^⊤^ follows a multivariate normal distribution 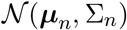 with mean vector ***μ***_*n*_ = (*μ*(**z**_1_),…, *μ*(**z**_n_))^⊤^ and covariance matrix 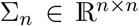, where *Σ_n,ij_* = *κ*(**z**_i_, **z**_j_) for *i*, *j* = 1,…, *n*. For a given set of cost function evaluations 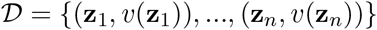, the structure of the Gaussian process allows the formulation of the conditioned surrogate model

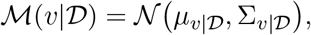

where the components of this distribution are

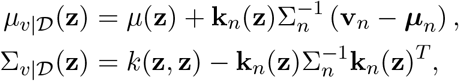

with **k**_n_(**z**) = (*κ*(**z**, **z**_1_),…, *κ*(**z**, **z**_n_)).

The first component of the weighted sum in (3.2) corresponds to the exploitation of low mean values and the second component incorporates the exploration of regions of the parameter space with high uncertainty regarding the value of the cost function *υ*. It is common to add a tuning parameter *ξ* into the acquisition function so that

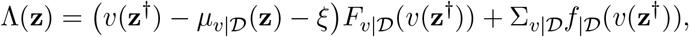

and increasing values of *ξ* lead to increased exploration by the algorithm. In our work we use *ξ* = 0.01, which is the common default value (Brochu et al., 2010). For the mean function *μ* in (3.1) we follow the usual convention of *μ*(**z**) ≡ 0 and for the covariance kernel *κ* we choose the popular Matérn function with *ν* = 1/2, see Rasmussen (2003) for information on many of the available alternatives.

### 3.3 Pipeline to simulate root-induced preferential flow through vegetated soil

A computational pipeline was assembled to generate simulations of moisture transport through vegetated soil from architecture data for the incumbent root system, illustrated by the green arrows in Figure 2. The pipeline takes as input a list of data which explicitly states the position, size and orientation of each segment in the root system. This is available for some experimental studies of plant roots, see e.g. (Danjon et al., 1999; Dupuy et al., 2005), as output of software packages that model the growth of root systems, e.g. CRootBox (Schnepf et al., 2018) and OpenSimRoot (Postma et al., 2017), or from simulation algorithms such as Algorithm 1 in Section 2. The pipeline then employs the kernel-based method presented in Section 2 to construct the volumetric density distribution and heterogeneity matrix for the root system, and subsequently parametrise the facilitation model (2.12), (2.16). The evolution of pore-water pressure over time through the vegetated soil is predicted by simulating, using the finite-element scheme, solutions to the facilitation model given the initial pressure distribution, the moisture infiltration conditions and the hydraulic properties of the soil. By employing Algorithm 2 to minimise the required cost function, see Section 3.4, the pipeline optimally calibrates the facilitation model (2.12), (2.16) with respect to the experimental data for the saturated hydraulic conductivity of the soil that is occupied by the root system in question, see e.g. Table 1. The parameter values which minimise the cost function and the numerical solutions to the corresponding optimal parametrisation of (2.12), (2.16) are the output of the pipeline.

**Figure 2:**
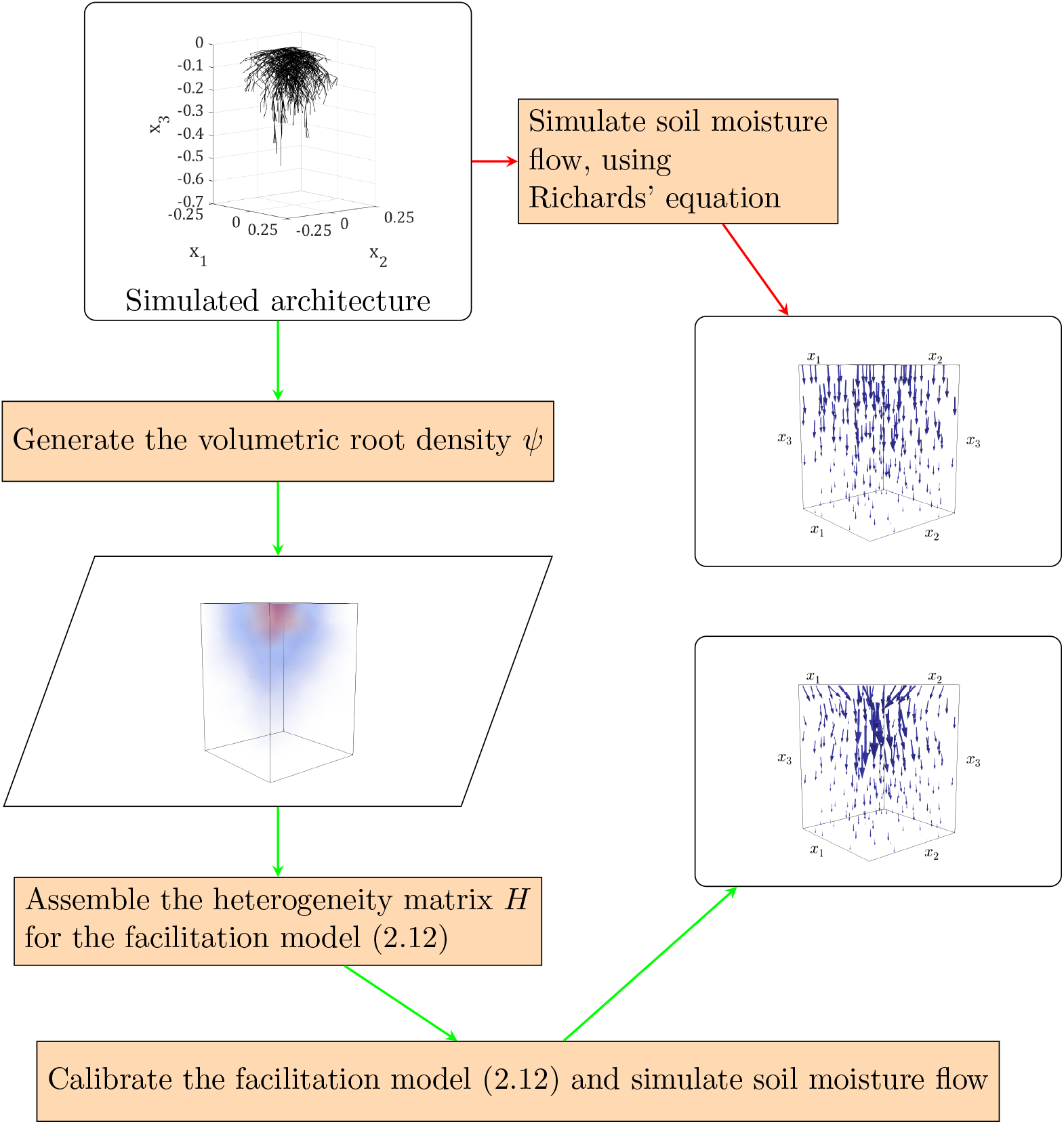
An illustration of the pipeline which calibrates the facilitation model (2.12), (2.16) for a given root system. Red arrows indicate the pipeline of the standard Richard’s equation (Richards, 1931), where no influence of the root system on soil-moisture transport is incorporated. Green arrows indicate the new pipeline of the facilitation model (2.12), (2.16), with which the preferential transport of soil moisture along root axes can be incorporated and its influence analysed.

The implementations of the finite-element scheme were carried out using the FEniCS library (Alnæs et al., 2015) for Python 3. Applications of Algorithm 1 to simulate root architecture data and implementations of the kernel-based methods for constructing volumetric root density distributions and heterogeneity matrices were carried out using the NumPy and SciPy libraries (Harris et al., 2020; Virtanen et al., 2020). For the implementation of Bayesian optimisation, to minimise the corresponding cost functions, the scikit-optimise library for Python 3, designed by Head et al. (2018), was used, whereas the application of Powell’s method was carried out using the *optimize* module of the SciPy library. Numerically computed quantities such as pore-water pressure, pressure head and soil-moisture profiles are visualised using the Paraview software (Ahrens et al., 2005), to show the influence of preferential moisture transport along the axes of a given root system on the moisture distribution in vegetated soil.

### 3.4 Numerical experiments to test hypotheses on the hydraulic properties of vegetated soil

We used the facilitation model (2.12), (2.16) to test the following hypotheses:

Q1 Is our optimisation scheme capable of effectively estimating

i. the facilitation constant *c_a_* which optimally calibrates the facilitation model (2.12), (2.16) when parametrising for a specific simulated root system?
ii. the facilitation constant *c_a_* and the parameters 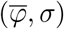 in Algorithm 1 for simulation of a truncated normal root system, which optimally parametrise and calibrate the facilitation model (2.12), (2.16)?
Q2 Do solutions of the facilitation model (2.12), (2.16) support the hypothesis that an increase in root abundance within soil leads to an increase in the infiltration rate of moisture through the soil?
Q3

i. When incorporating information on the structure of a root system, as well as root abundance, do simulated profiles from the facilitation model (2.12), (2.16) support the hypothesis that the transport of moisture along root axes significantly influences soil hydraulic properties?
ii. Is it possible to effectively calibrate parametrisations of (2.12), (2.16), which incorporate the structure of root systems of different plant species, e.g. grass and willow plants, by using the same facilitation constant ca for both plant species?

Questions Q1, Q2 and Q3 are addressed by our results in Sections 4.1, 4.2 and 4.3 respectively. The remainder of this section specifies the cost functions that we minimise to calibrate the facilitation model (2.12), (2.16) in an attempt to answer questions Q1-Q3.

To calibrate (2.12), (2.16) with respect to the experimental data, the first step is to parametrise the empirical model (2.13), (2.16) using the 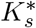 values in Table 1 and then solve numerically to obtain the pressure head *h*_e_j__, where the subscript j indicates whether a grass (j = g) or willow (j = w) root system is considered. These empirical pressure head solutions provide the experimental benchmark against which solutions from the facilitation model (2.12), (2.16) can be judged. Given the empirical pressure head profiles *h*_e_j__ corresponding to 8 week old grass and willow plants, the corresponding laterally averaged pore-water pressure profiles 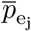 (kPa) are computed as

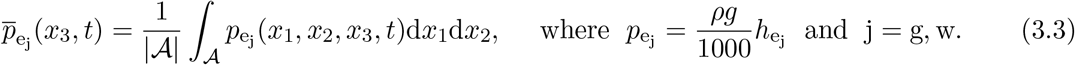

To address Q1(i), the first step is to construct the volumetric root density function *ψ* which corresponds to the specific simulated root system. Given an axial facilitation constant *c*_a_j__ > 1, we are then able to construct the heterogeneity matrix *H*(*c*_a_j__; *x*) and hence parametrise (2.12), (2.16) with respect to this root system. Solutions to this parametrisation of (2.12), (2.16) are then obtained numerically and, using the same formula as in (3.3), the laterally averaged pore-water pressure 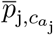 is computed. The optimal *c*_a_j__ > 1 is then found by minimising the cost function

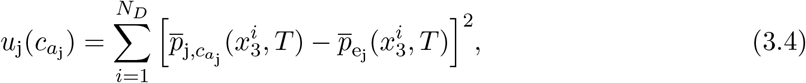

where *T* > 0 is the final time and *N_D_* > 0 is a number of points 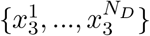 on the interval [*D*, 0].

Testing Q1 (ii) involves determining the optimal 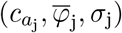 with which to simulate a truncated normal root system of species j, and then parametrise the corresponding facilitation model (2.12), (2.16). Given parameters 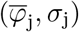 for root species j, we use Algorithm 1 to construct a truncated normal root system. With this root system and an axial facilitation constant *c*_a_j__, the corresponding density function *ψ* and heterogeneity matrix *H*(*c*_a_j__; *x*) are constructed and a full parametrisation of (2.12), (2.16) is formulated. The laterally averaged pore-water pressure from this parametrisation is denoted as 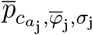. Then the optimal parameters 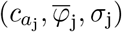 are found by minimising the cost function

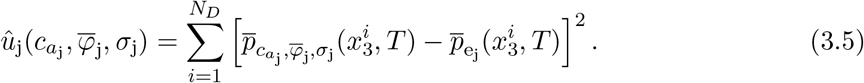

We address Q2 by using Algorithm 1 to simulate 8 week old uniform grass and willow root systems. Using the kernel-based method described in Section 2, the corresponding volumetric density functions and heterogeneity matrices are constructed to obtain parametrisations of (2.12), (2.16) for both uniform root systems. The cost function *u*_j_ in (3.4) for each uniform root system is then minimised to find the optimal axial facilitation constant *c*_a_j__. The simulated profiles from optimal parametrisations of the facilitation model for both root systems are then investigated to determine if root abundance is increasing infiltration. Since, in these root systems, the orientation of root segments depends upon parameters which are all drawn from uniform distributions, the model solutions do not provide information on the impact of the structure of the root system on infiltration rate, instead they only indicate the influence of root abundance on the moisture distribution in soil.

Question Q3 (i) is tested by using Algorithm 1 to simulate 8 week old truncated normal grass and willow root systems. Since the grass root system is deemed to have roots which are oriented more horizontally we simulate it using parameters 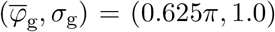. The willow root system is deemed to have roots which are oriented more vertically and we simulate it using parameters 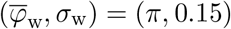. We again construct the volumetric density function and heterogeneity matrix for each system to obtain parametrisations of (2.12), (2.16). The cost function *u*_j_ in (3.4) corresponding to the parametrisation for each truncated normal root system is then minimised to find the optimal axial facilitation constant *c*_a_j__, for j = g, w. Since the structures of the truncated normal root systems are no longer uniform, the corresponding parametrisations of (2.12), (2.16) truly incorporate the influence of preferential soil moisture transport along root-axes within a given system. If the simulated profiles from the optimal parametrisations of (2.12), (2.16) agree more closely to the corresponding empirical profiles than when parametrising with respect to uniform root systems, then this will provide evidence that the transport of soil moisture along root axes is a determining factor in how the hydraulic properties of soil are influenced by an incumbent root system.

Question Q3 (ii) is first tested by taking the truncated normal willow and grass root systems simulated to test Q3 (i) along with the corresponding volumetric density functions, heterogeneity matrices and parametrisations of the facilitation model (2.12), (2.16). Then, considering a single axial facilitation constant *c_a_*, i.e. *c_a_g__* = *c_a_* and *c_a_w__* = *c_a_*, we minimise the cost function

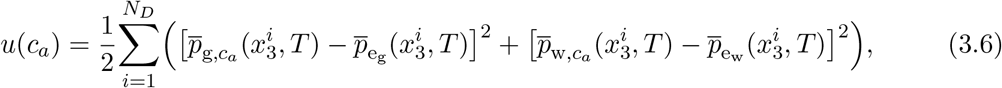

to find a cross-species optimal facilitation constant *c_a_*. If pore-water pressure profiles from the parametrisations of (2.12), (2.16), for these truncated normal willow and grass root systems, which use the same *c_a_* value, can achieve a better fit to the empirical profiles than those which come from uniform root systems then this will suggest that it is the structure of the root systems that is causing the strong agreement with experimental data. This will therefore strengthen the evidence that preferential soil moisture transport along root axes is a determining factor in how the hydraulic properties of vegetated soils are influenced by incumbent root systems. To provide further evidence to support this hypothesis we minimise the cost function

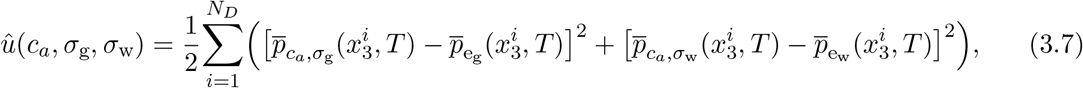

to determine an optimal (*c_a_*, *σ_g_*, *σ_w_*). Here (0.625π, *σ_g_*) and (π, *σ*_w_) are the parameters used to simulate truncated normal root systems of grass and willow respectively and *c_a_* is the single crossspecies axial facilitation constant that is used to complete the parametrisation of (2.12), (2.16) for each of the simulated truncated normal root systems. If pore-water pressure profiles from parametrisations of (2.12) (2.16), for these root systems generated with the optimal (*σ_g_*, *σ_w_*), which use the same *c_a_* value, can achieve a better fit to the experimental data in Table 1, then this will further strengthen the evidence that preferential soil moisture transport along root axes is a determining factor in how the hydraulic properties of vegetated soils are influenced by an incumbent root system.

## 4 Results

### 4.1 Estimation of parameter values for the facilitation model

First, we tested whether the optimisation scheme can successfully estimate an optimal value for the facilitation coefficient 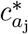 given empirical data, i.e. Q1 (i) from Section 3.4. We minimised cost function *u*_j_, defined in (3.4), to find 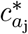 which optimally parametrised the facilitation model (2.12), (2.16) for a specific truncated normal grass root system simulated using parameter values 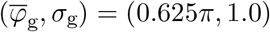. In this scenario the use of Bayesian optimisation provided a moderate advantage over Powell’s method (Table 2). For this specific grass root system, Powell’s method was able to locate a value 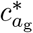 with small value for 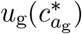. However, Bayesian optimisation located a very similar optimal value 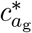, with equally low 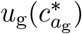, in approximately half the time taken by Powell’s method, see Table 2.

**Table 2:** Convergence results for Powell’s method and Bayesian optimisation. All schemes were seeking the minimiser 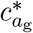 of the one-dimensional cost function *u*_g_, given by (3.4), which optimally calibrated the facilitation model (2.12), (2.16) with respect to a specific truncated normal grass root system, simulated using the parameters 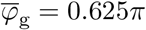 and *σ*_g_ = 1.0.

| Method | Bayesian Optimisation |  |  |  | Powell's |
| --- | --- | --- | --- | --- | --- |
| Function evaluations | 5 | 10 | 15 | 20 | 20 |
| Run times, (hr,min,sec) | 44m 23s | 1h 28m 41s | 2h 13m 33s | 3h 3m 45s | 2h 58m 51s |
| $c_{a_g}^*$ | 668.66 | 669.39 | 669.15 | 668.95 | 669.44 |
| $u_g(c_{a_g}^*)$ | 0.05083 | 0.05077 | 0.05078 | 0.05079 | 0.05077 |

To test whether the optimisation scheme can successfully estimate both the value of the facilitation coefficient and the distribution of the root angles from the positive vertical axis, given experimental data i.e. Q1 (ii), we minimised the cost function 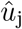, given by (3.5). This identified the optimal parameters *c*_a_g__ and 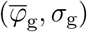, with which to simulate a truncated normal grass root system and parametrise (2.12), (2.16) (Table 3). Due to the use of random number generators in Algorithm 1, two root systems that are simulated using the same 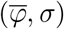 will never have exactly the same structure. This adds noise to the optimisation process, and, together with the increase in the parameter space dimension from one to three, makes the existence of multiple minimisers likely. As a result Powell’s method performed significantly less efficiently than for the one-dimensional optimisation problem (Tables 2 and 3). Bayesian optimisation however, did not suffer the same deterioration in performance when the dimensionality of the problem was increased. It can be seen in Table 3 that a Bayesian optimisation scheme with 20 function evaluations identifies parameter values that minimise 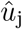 to the same extent as the optimum identified by Powell’s method after one iteration (64 function evaluations). However, Bayesian optimisation achieves this result in less than one-third of the computation time taken by Powell’s method.

**Table 3:** Convergence results for optimisation schemes which seek minimisers of the cost function 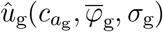, see (3.5), when considering simulated 8 week old truncated normal grass root systems. The minimiser was the parameter triple 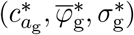, such that simulating a truncated normal 8 week old grass root system with parameters 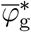 and 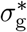, then generating *ψ*(*x*) and 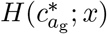 for this simulated system, resulted in the facilitation model (2.12), (2.16) being optimally calibrated to the data in Table 1. The parameter space from which we sought a minimiser was 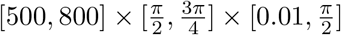 and the randomly chosen start point used for each scheme was 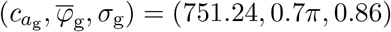.

| Method | Bayesian Optimisation |  |  | Powell's |
| --- | --- | --- | --- | --- |
| Function evaluations | 5 | 10 | 20 | 64 |
| Run times, (hr, min, sec) | 50m 26s | 1h 44m 32s | 3h 28m 42s | 12h 15m 8s |
| $c_{a_g}^*$ | 640.44 | 730.54 | 561.83 | 612.15 |
| $(\bar{\varphi}_g^*, \sigma_g^*)$ | $(0.70\pi, 0.96)$ | $(0.57\pi, 0.84)$ | $(0.74\pi, 1.48)$ | $(0.74\pi, 0.48)$ |
| $\hat{u}_g(c_{a_g}^*, \bar{\varphi}_g^*, \sigma_g^*)$ | 0.27 | 0.17 | 0.11 | 0.11 |

Both results from this section indicate that our calibration pipeline is effective in determining parametrisations of model (2.12) (2.16) which produce pore-water pressure profiles that strongly agree with the experimental results.

### 4.2 An increase in root abundance causes an increase in infiltration rate

To test the hypothesis that an increase in root abundance within soil leads to an increase in the infiltration rate (question Q2) the cost functions *u*_g_ and *u*_w_, see (3.4), were minimised to obtain parametrisations of facilitation model (2.12), (2.16) for specific grass and willow root systems in which roots were not assumed to grow at any preferred angle, see Algorithm 1. The results in Figure 3 and columns 2 and 5 of Table 4 show that laterally averaged pore-water pressure profiles from the optimal parametrisations of (2.12), (2.16) that corresponded to these uniform root systems, agree strongly with the corresponding empirical pore-water pressure profiles based on the experimental data of Leung et al. (2018). This again shows that the calibration pipeline is effective and also supports the observation of Leung et al. (2018) that root systems induce increased infiltration (reduction in pore-water pressure) in the top soil where volumetric root density is greatest. Furthermore, Figure 3 also shows that the minimum pore-water pressure is lower in soil vegetated by willow than by grass. Thus the optimally parametrised facilitation model (2.12) (2.16), for soils vegetated by simulated uniform root systems, predicts that more infiltration will occur if soil is vegetated by willow than by grass.

**Figure 3:**
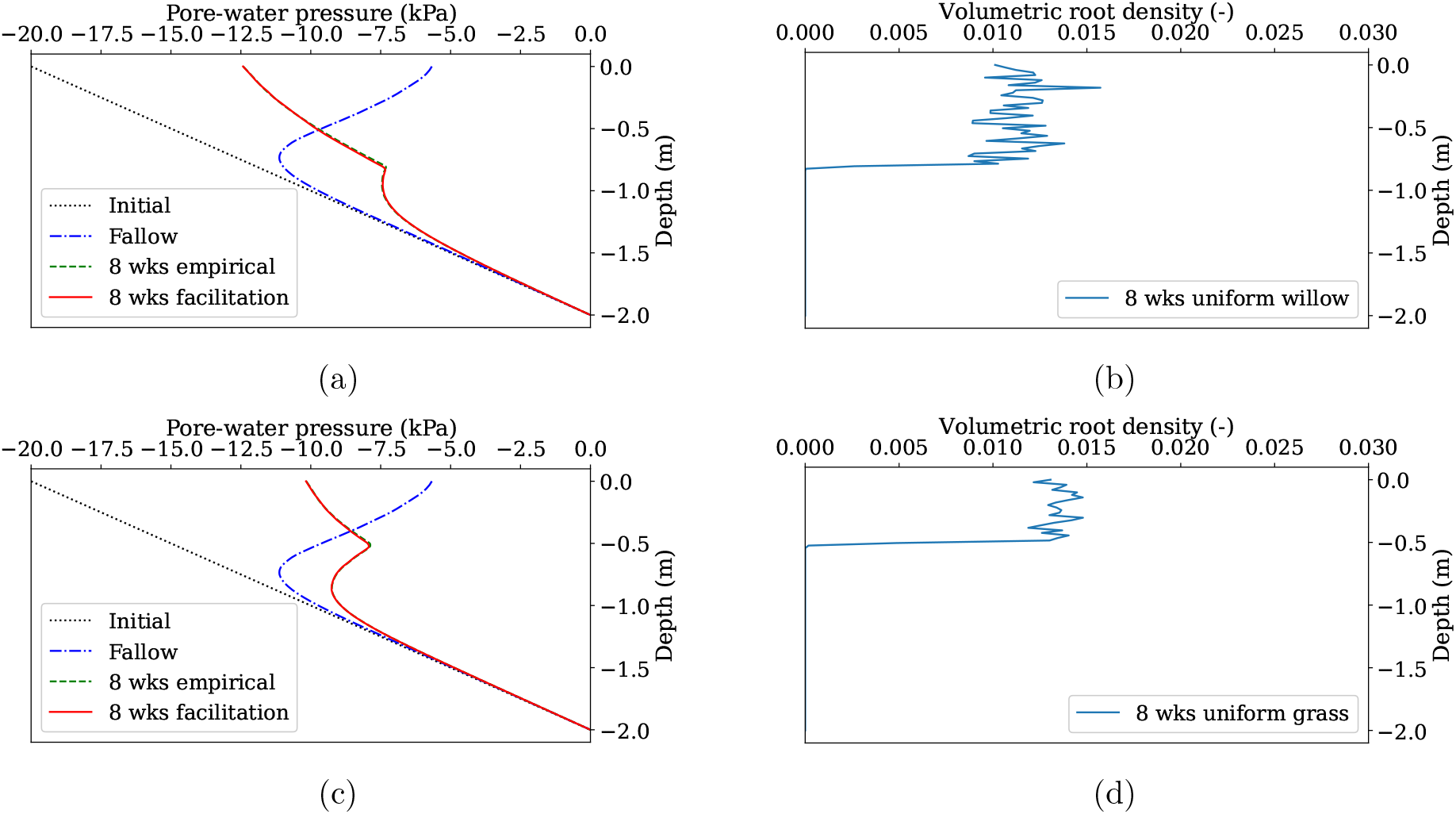
Plots (a) (willow) and (c) (grass) show the laterally averaged pore-water pressure profiles, after 2 days of rainfall, simulated from the empirical model (2.13), (2.16) and from optimal parametrisations of the facilitation model (2.12), (2.16) for simulated uniform root systems. The values of the axial facilitation constant *c*_a_j__, with j = g, w, that optimally parametrise (2.12), (2.16) for the simulated uniform root systems are shown in Table 4. Plots (b) and (d) show the laterally averaged profile of the volumetric root density *ψ* that was used to obtain the parametrisation of (2.12), (2.16) for each simulated uniform root system.

**Table 4:** The axial facilitation constants 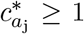 (j = w for willow and j = g for grass) which optimally calibrate model (2.12), (2.16) for simulated 8 week old willow and grass root systems. The optimal values were found by minimising the cost function *u*_j_ for each of the 4 root system types, see (3.4) for the definition of *u*_j_.

| Species | Grass |  | Willow |  |
| --- | --- | --- | --- | --- |
| Distribution of root polar angles $\varphi_i$ over $[\frac{\pi}{2}, \pi]$, in the simulated root system | Uniform | Truncated Normal<br>$\bar{\varphi}_g = 0.625\pi$<br>$\sigma_g = 1.0$ | Truncated Normal<br>$\bar{\varphi}_w = \pi$<br>$\sigma_w = 0.15$ | Uniform |
| $c_{a_j}^*$ | 597.49 | 669.25 | 662.53 | 1254.04 |
| $u_j(c_{a_j}^*)$ | 0.0845 | 0.0508 | 0.0913 | 0.3143 |

Analysis of the parameters obtained by optimisation, also revealed that drastically different optimal axial facilitation constants *c_a_g__* and *c_a_w__* were obtained when calibration of the facilitation model was performed for the uniform willow and grass root systems. This result indicates that while these parametrisations of the facilitation model agree well with the experimental data in Table 1, they do not reflect any influence of preferential moisture transport along the axes of roots in differently structured willow and grass root systems.

### 4.3 Predictions of soil moisture transport are improved by incorporating structure of root systems

We also tested whether numerical simulations for facilitation model (2.12), (2.16) support the hypothesis that the transport of moisture along root axes significantly influences soil hydraulic properties (Q3 (i)). This was done by minimising cost functions *u*_g_ and *u*_w_ to find the optimal values of 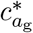 and 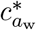 when considering grass and willow root systems that were simulated using parameters 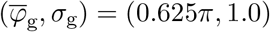 and 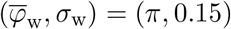 respectively (Table 4). Columns 3 and 4 of Table 4 show that the calibration pipeline is effective for truncated normal root systems and that the laterally averaged pore-water pressure profiles simulated from (2.12), (2.16) agreed strongly with the empirical profiles simulated from (2.13), (2.16). The squared Euclidean distances 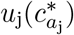 between the pressure profiles simulated from the facilitation model (2.12), (2.16) and the corresponding profiles from (2.13), (2.16) are also reduced when considering parametrisations for truncated normal root systems instead of uniform (Table 4). Furthermore, the optimal facilitation constants corresponding to the truncated normal root systems for both species are significantly more similar than those corresponding to the uniform root systems.

Finally, we tested whether a single facilitation coefficient can accurately describe the influence of both willow and grass root systems on soil moisture transport (Q3(ii)). Results which support this hypothesis would indicate that the different influences on soil moisture transport, that are induced by the root systems of different plant species, could be explained by one global mechanism. The cost function *u*, defined in (3.6), was minimised to investigate whether it is possible to complete the parametrisations of (2.12), (2.16) that correspond to truncated normal willow and grass root systems, simulated using 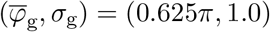 and 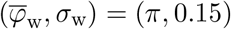 respectively (Table 5), by using a single common facilitation coefficient. Results show that this is indeed possible (Table 5 and Figures 4 (a) and (c)) and that pore-water pressure profiles from these parametrisations of (2.12), (2.16) for the truncated normal root systems, which use a cross-species optimal value 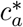, still agree more strongly with the experimental data than profiles from optimal parametrisations of the facilitation model for simulated uniform root systems. Figures 4 (f) and (h) show that parametrisations of the facilitation model for the truncated normal willow and grass root systems, which have the same value of facilitation constant 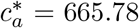, produce pore-water pressure profiles for which the squared Euclidean distance from the empirical profiles is less than when parametrising for uniform root systems using the same value for 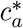. Furthermore, Figures 4 (e) and (g) show that differences in the profiles of the soil pore-water pressure produced by the parametrisations of (2.12), (2.16) for these uniform and truncated normal root systems occur at the shallow soil depths where root density is highest. These results indicate that incorporating information on root system structure into (2.12), (2.16) leads to the improved agreement of model solutions with experimental results, whilst also providing evidence of the existence of an optimal cross-species facilitation constant 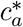. Since (2.12), (2.16) incorpo-rates explicit root structure information by simulating preferential soil moisture transport along the axes of the structure, these results provide evidence that the preferential transport of soil moisture along root axes is a determining factor in how root systems influence the hydraulic properties of soil. To further support this, the results in Table 6 show that it is possible to estimate the optimal parameters 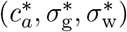, with a cross-species axial facilitation constant 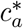, where truncated normal willow and grass root systems are constructed using parameters 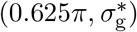 and 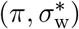.

**Figure 4:**
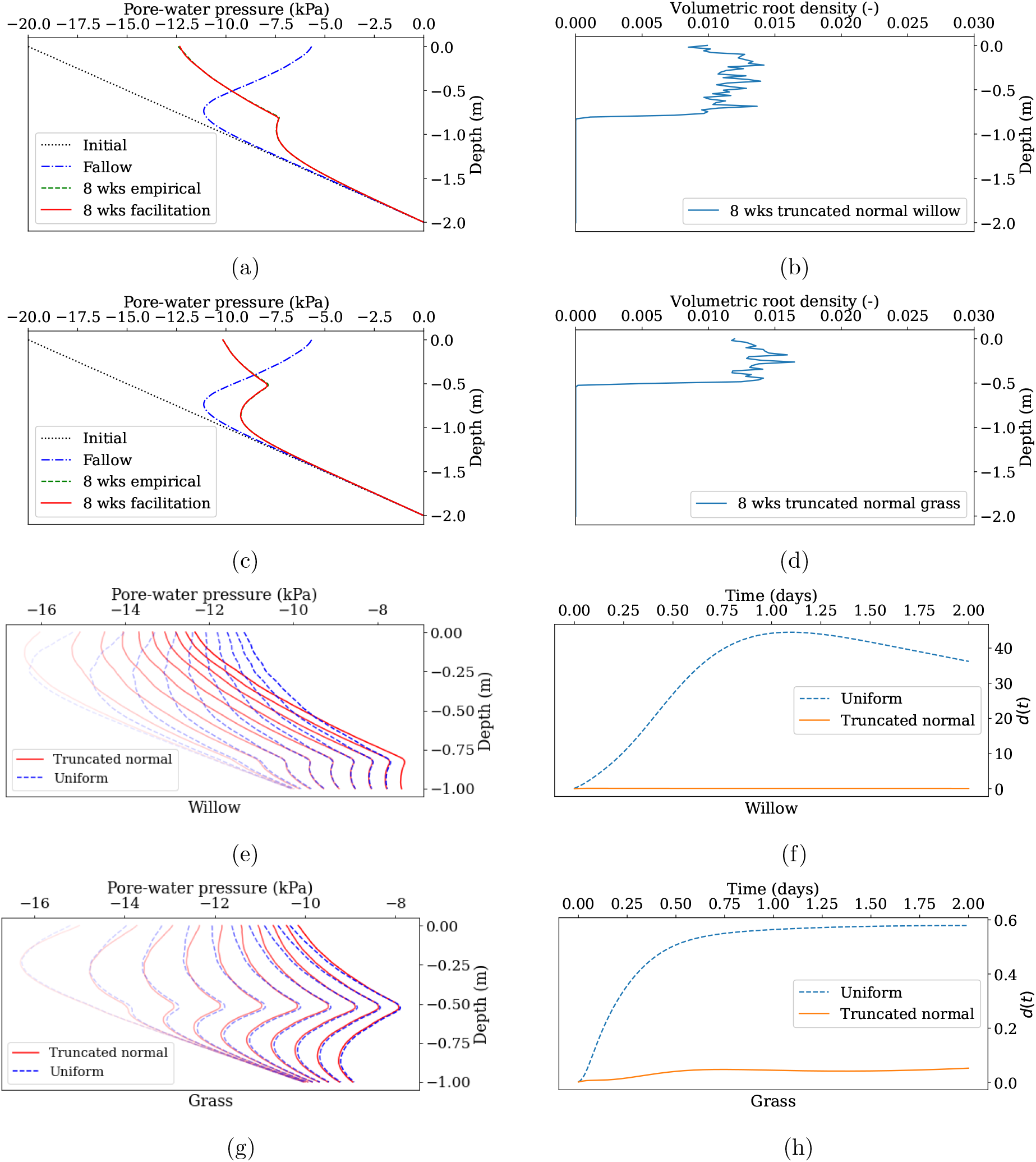
Plots (a) (willow) and (c) (grass) show laterally averaged pore-water pressure profiles after 2 days of rainfall, simulated from (2.13), (2.16) and from optimal parametrisations of (2.12), (2.16) for truncated normal root systems with parameters 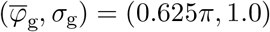 for grass and 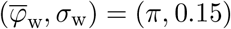 for willow. Plots (b) and (d) show the laterally averaged profiles of the corresponding volumetric root densities. Plots (e) and (g) show laterally averaged porewater pressure profiles simulated from parametrisations of (2.12), (2.16) for each of the simulated root systems at different times. The faintest and leftmost lines correspond to earliest simulation times and the lines of full opacity (rightmost) show the final profiles. Plots (f) and (h) show the evolution over time of the squared Euclidean distance 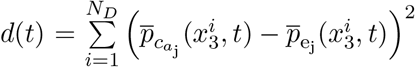, between the laterally averaged pressure profiles from (2.12), (2.16) and (2.13), (2.16), see (3.3) for the definitions of 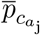 and 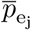. In all the results in this figure, the facilitation model (2.12) (2.16) was parametrised for truncated normal and uniform root systems of both grass and willow using the cross-species optimal 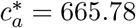 from Table 5.

**Table 5:** The optimal cross-species axial facilitation constant 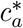 which minimises *u*, see (3.6). Using this value, the facilitation model is calibrated effectively for soils vegetated by truncated normal root systems of either grass or willow, simulated using the parameter values in row 2.

| Species | Grass | Willow |
| --- | --- | --- |
| Parameters in truncated normal distribution for $\varphi_i$ over $[\frac{\pi}{2}, \pi]$ | $(\bar{\varphi}_g, \sigma_g) = (0.625\pi, 1.0)$ | $(\bar{\varphi}_w, \sigma_w) = (\pi, 0.15)$ |
| Optimal cross-species $c_a^*$. | 665.78 | |
| $u(c_a^*)$ | 0.0723 | |
| $u_j(c_a^*)$ | 0.0521 | 0.0925 |

**Table 6:** The optimal (*c_a_*, *σ*_g_, *σ*_w_) found by Bayesian optimisation to minimise 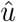, where 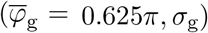 and 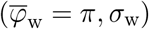 are the parameters of the truncated normal distributions that are used to simulate the grass and willow root systems respectively and *c_a_* is the cross-species axial facilitation constant used in the corresponding parametrisations of the facilitation model. The scheme searches for the optimal *σ*_g_ and *σ*_w_ within the interval 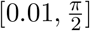 and the optimal *c_a_* within the interval [550, 750].

| Function evaluations | Random start point $(c_a, \sigma_g, \sigma_w)$ | Optimum $(c_a^*, \sigma_g^*, \sigma_w^*)$ | $\hat{u}(c_a, \sigma_g, \sigma_w)$ |
| --- | --- | --- | --- |
| 25 | (683.80, 0.28, 0.71) | (693.41, 0.82, 0.11) | 0.15 |

## 5 Discussion

### 5.1 Improving models by incorporating preferential soil moisture transport along root axes

The influence of root systems on soil hydraulics is modelled in many diverse ways. Approaches often involve modifying Richards’ equation, for example by introducing a sink term for water uptake that depends on the root biomass distribution, see e.g. Leung et al. (2015), Wu et al. (1999). Such modifications are simple and effective, however they simplify greatly the effect of the root system on the moisture distribution in soil, since properties like root diameter distribution and branching angle are not well represented.

Richards’ equation for moisture transport in the soil, coupled with the Darcy flux for axial moisture transport through root tissue, was used to solve simultaneously for moisture transport in the soil and in the root system, see e.g. Amenu and Kumar (2008), Arbogast et al. (1993), Doussan et al. (2006). The coupling is often used to simulate water uptake by plants and is achieved by adding a sink/source term to each equation which is proportional to the radial conductivity of the root tissue and the difference in matric potential between the soil and root moisture. The influence of root abundance is incorporated through the dependence of the radial hydraulic conductivity of root tissue on the fraction of roots which occupy that layer of soil, however no dependence of the soil’s hydraulic conductivity on root abundance or root system structure is considered, despite the array of experimental evidence supporting the existence of such a relationship, see e.g Wilcox et al. (2003), Scholl et al. (2014), Song et al. (2017), Leung et al. (2018). It is possible to envisage modification of these models to include both water uptake and facilitation of water flow along root axes. However, these models rely on computationally intensive simulations, resolving individual branches of a root system, which prohibit their application to soil volumes larger than those occupied by a few plants.

Dual-porosity and dual-permeability models are a popular approach for incorporating, into simulations of soil moisture transport, the water flow along soil fractures, cracks or channels that are created by plant roots see e.g. Gerke and Van Genuchten (1993), Granier et al. (1999), Simunek et al. (2003), Simunek et al. (2005), van der Heijden et al. (2013), Valle et al. (2017). These models either employ a coupling of different parametrisations of Richards’ equation to model the moisture transport in the fractures and the soil matrix, or they couple a transport equation or ordinary differential equation for the fracture flow with Richards’ equation for the moisture transport in the soil matrix. The coupling in both approaches is achieved using a water transfer rate. Rigorous derivation of dual-porosity models from the microscopic description of the two pore system structure of soil can also be obtained via homogenization techniques, see e.g. Arbogast et al. (1990), Bourgeat et al. (2003), Bourgeat et al. (1996), Hornung (1996). Despite their ability to model flow along soil fractures and channels, dual-porosity/dual-permeability models do not impose any dependence upon the structure of the root systems that vegetate the soil, even though this has a likely influence on the orientation and size of preferential flow channels (Noguchi et al., 1999).

In this work we addressed the shortcomings of the existing models for moisture transport in vegetated soil by modifying Richards’ equation to formulate the facilitation equation (2.12). Our model introduces the preferential transport of soil moisture along root axes by using a volumetric root density function *ψ* and a heterogeneity matrix *H* to impose a dependence of the soil moisture flux on root abundance and orientation. In regions of high volumetric root density the facilitation model promoted local preferential moisture transport parallel to the axes of roots in that section of the underlying system. With comparison to previous studies, our approach has addressed two main limitations of existing models. Firstly, the incorporation of root architectural data into the heterogeneity matrix allows us to obtain numerical simulations of soil hydraulic processes which account for the influence of the complex structure of the incumbent root system. This is lacking in previous approaches such as dual-porosity/dual-permeability models. Second, since the root system structure is aggregated in the heterogeneity matrix, explicit representation of root systems is not needed and performing simulations at a wide range of spacial and temporal scales is possible. This overcomes the limitation of approaches where a root branching network needs to be simulated explicitly. Clearly, the incorporation of root architectural properties into the heterogeneity matrix does not consider explicitly the moisture flow around a single root, which may be a limitation for some applications.

We tested the facilitation model against the data of Leung et al. (2018) for the saturated hydraulic conductivity of soils vegetated by willow and grass. This involved simulating the willow and grass root systems, see Algorithm 1, and applying our calibration pipeline to parametrise the facilitation model for each of the systems. Results showed that for simulated truncated normal root systems of 8 week old willow and grass, with parameter values chosen to reflect the data of Leung et al. (2018), the optimal c_aj_ values converged and the pressure profiles from the optimally parametrised facilitation model agreed more strongly with the experimental data than in the case of uniform root systems (Figures 3, 4 and Table 4). This therefore showed that the main cause of the difference between solution profiles from parametrisations of the facilitation model was the data-driven heterogeneity that had been imposed on the structure of the simulated truncated normal root systems, and the preferential flow that this induced. Furthermore, it suggested that the capacity of the facilitation model to incorporate the preferential flow along the axes of roots, was what caused the agreement with the experimental data to improve when considering the truncated normal root systems.

### 5.2 Acquiring data with which to parametrise the facilitation model

With our new model (2.12), (2.16), we have been able to develop a pipeline which goes from the architecture data for a given root system to the simulated profile of the preferential moisture transport that it induces (Figure 2). The assembly of this pipeline requires the decomposition of the given root system into components, i.e. conical frustums, where the length along with the centre points and diameters of the proximal and distal ends are available for each frustum. Such data have been obtained previously by Danjon et al. (1999) who analysed excavated root systems. More recently, x-ray computed tomography has proved a viable method of imaging the 3D architecture of a root system without the need for excavation (Mairhofer et al., 2013; Zhao et al., 2020). The process time and accuracy of x-ray computed root system reconstructions is improving significantly (Gao et al., 2019), and applying the analysis techniques to these threedimensional images could provide an efficient and non-invasive method of obtaining the root architecture data required for our model (2.12), (2.16). Structural data for particular species of root systems could also be estimated by making use of the numerous software tools that model the growth of root systems and output the corresponding architecture data, see for example CRootBox (Schnepf et al., 2018), OpenSimRoot (Postma et al., 2017) and DigR (Barczi et al., 2018). A more extensive list of architecture simulation tools can be found in Passot et al. (2019) along with a review of the computation pipelines into which they could potentially be integrated. The volumetric root density function *ψ* needed to compute heterogeneity matrices can more readily be obtained from solutions to partial differential equations for root density used to model root system growth (Bastian et al., 2008; Dupuy et al., 2005, 2010). Solutions to these models are also more amenable to applications at different spatial and temporal scales, however, more work is needed to determine effective strategies for parametrising density based models with empirical data. In order to calibrate (2.12), (2.16) for a given root system, the effective saturated hydraulic conductivity of the soil that it occupies is also required. Provided those data on root system architecture and effective soil saturated hydraulic conductivity are available, our pipeline (Figure 2) can be applied to any root system and the influence of the root system on soil moisture transport can be analysed. Numerical simulations for (2.12), (2.16) carried out using architectural data for the root system of mature maritime pine, from Dupuy et al. (2005), demonstrate that the approach is suitable to complex simulation scenarios (Figure 5).

**Figure 5:**
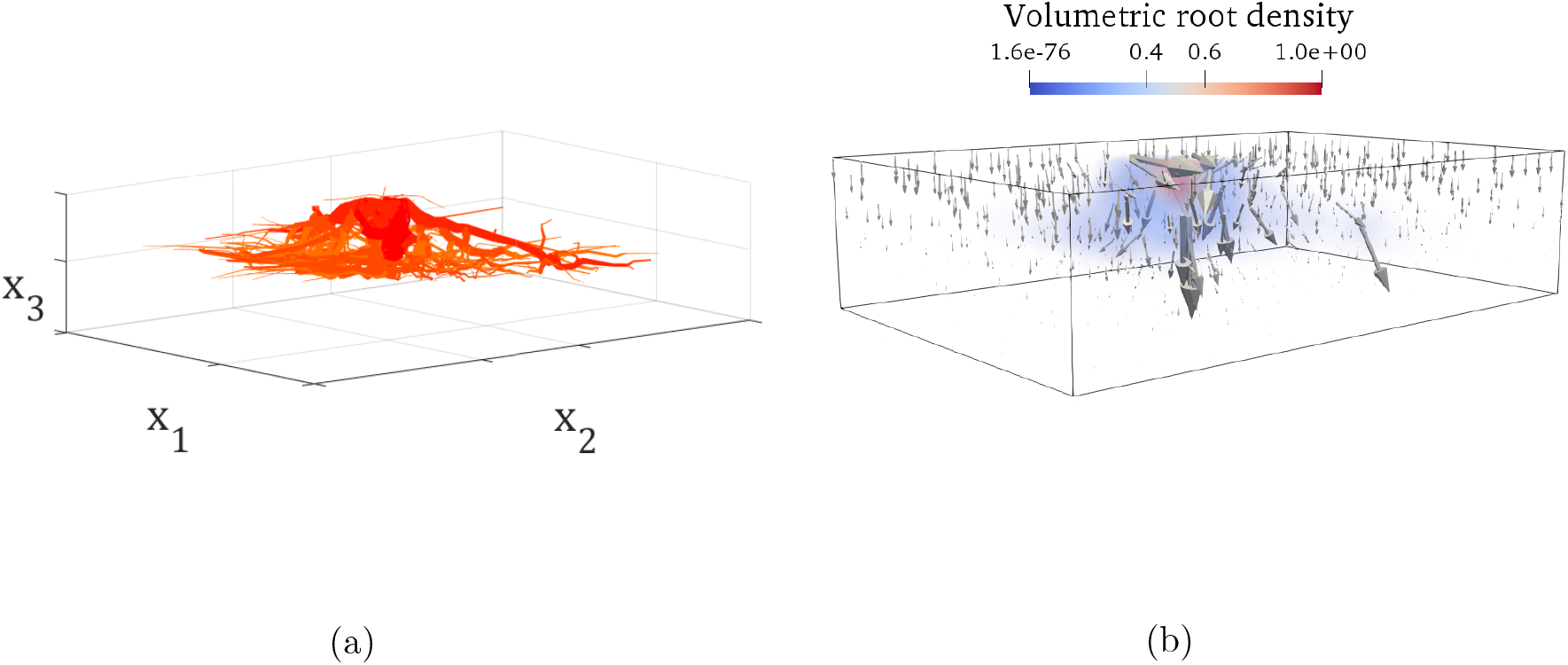
Figure (a) displays architecture data for a real pine root system that was studied by Dupuy et al. (2005) and expresses the root system as a union of conical frustums that occupy a soil domain. Figure (b) shows the volumetric density function *ψ* that was generated for the architecture in (a), using our methods from Section 2, and the resultant moisture transport from the parametrised facilitation model (2.12), (2.16). The direction of an arrow in (b) indicates the direction of flow and the size of an arrow indicates the magnitude of the flow.

### 5.3 Towards new approaches integrating the influence of vegetation into soil hydrological models

Vegetated soils are generally known to exhibit higher rates of moisture infiltration during rainfall and irrigation than bare soils (Huang et al., 2015; Leung et al., 2018). Below ground however, the impact of the preferential moisture transport caused by root systems is less clear. For example, results were obtained by Song et al. (2017) within the context of appropriate cover vegetation for landfill sites. They found that the preferential flow caused by Vetivier grass root systems facilitated downward moisture drainage through the landfill site, resulting in the leaching of harmful waste chemicals into water supplies. In contrast, Song et al. (2017) also reported that Bermuda grass root systems induced preferential flow within the soil which lead to reduced moisture drainage. Furthermore, when assessing the fallibility of a vegetated hill slope to landslide, it is not appropriate to simply assume that it will exhibit the same characteristics as a fallow hill slope (Leung et al., 2018). The experimental results of Lourenço et al. (2006) showed that hill slope regions where below ground pore-water pressure is high are more likely to become unstable, however it has been observed that these pore-water pressure distributions vary significantly depending on the vegetation cover of the hill slope (Ghestem et al., 2011).

Unfortunately, utilisation of vegetation within land management strategies, either as geotechnical or hydrologic barriers, is limited because it is difficult to predict the effect of vegetation types on the hydraulic properties of soil. Improved soil hydrological models could be used to inform the choice of vegetation cover so as to best manage soil-moisture flow patterns. For example, to inhibit excessive drainage from cultivated fields, or redistribute pore-water pressure during rainfall to maintain slope stability. Software packages such as TUFLOW (Syme, 2001) have been developed to model flood inundation and an overview of these is provided by Teng et al. (2017). The modelling in these software packages focuses largely on above ground water flow with some account taken for the different drying and wetting cycles of the affected soil. Incorporating a version of the facilitation model (2.12), (2.16) would enhance their ability to predict a soil’s buffer and infiltration capacity, given the root structure characteristics of the vegetation cover that is present.

Understanding the flow of water in agricultural soils is also essential to crop production. The hydraulic properties of a soil affect both water distributions and nutrient dynamics, highlighting the importance to optimise irrigation for water and nutrient uptake efficiency (van der Heijden et al., 2013). Industrial software packages, such as APSIM (Holzworth et al., 2014) and DSSAT (Jones et al., 2003), have been developed to predict crop responses to changing environmental conditions and both packages integrate groundwater flow models into their simulation pipelines. It is surprising, however, that none of these software account for preferential flow induced by the root systems of crops. The integration of facilitation models into crop modelling packages may enable targeted irrigation based on the size and architecture of the root systems of a given crop. This work shows that it is possible to account for the influence of root system architecture in a model for soil hydraulics. The approach developed here was calibrated and tested using simple experimental data. The challenge now is to confirm the results by considering a broader range of plant species and soil conditions. This would show that the model has the capacity to provide reliable strategic information to decision makers within land management.

## Acknowledgements

Andrew Mair was supported by The Maxwell Institute Graduate School in Analysis and its Applications, a Centre for Doctoral Training funded by the UK Engineering and Physical Sciences Research Council (grant EP/L016508/01), the Scottish Funding Council, Heriot-Watt University and the University of Edinburgh. We thank David Boldrin for fruitful discussions on root soil hydraulics.

## A Computation of a rotation matrix R for a particular root of a root system

For a component conical frustum 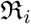 of the root system 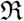, let **v** be the direction vector for 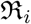. A rotation matrix 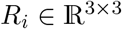 can be computed, which by left multiplication rotates the vector **e**_3_ = (0, 0, 1)^⊤^ into the direction of 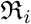. We compute *R_i_* for a given 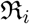 using the formula of Rodrigues (1840) as follows:

1. Normalised direction vector of the segment: 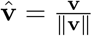.
2. Normalised cross product of the segment: 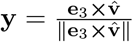.
3. Angle between **e**_3_ and 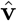 around the axis 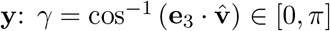.
4. Cross product matrix of **y**:

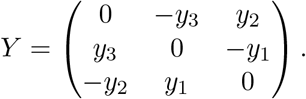
5. Rotation matrix:

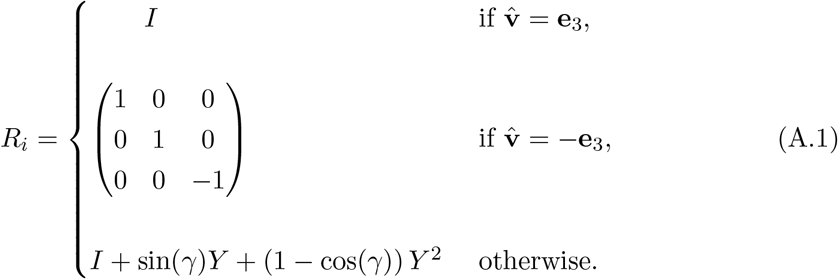

## B Properties of the heterogeneity matrix H

### Proposition B.1

*For any axial facilitation constant c_a_* ≥ 1, *the heterogeneity matrix H*(*c_a_*; *x*) *of any root system 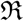 is uniformly positive-definite i.e, there exists* λ > 0 *such that*

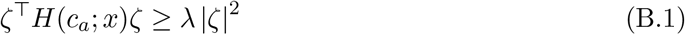

*for all x* ∈ Ω *and all* 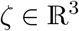.

*Proof*. The heterogeneity matrix has the form 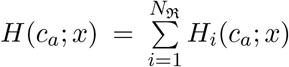 where, for each segment 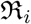 in the underlying root system, the associated matrix is 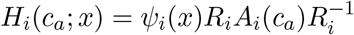. Given a segment 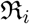, let *ϑ_i_* ∈ [0, 2π) be the anti-clockwise angle of the segment from the positive *x*_1_ axis and 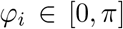 be the angle of 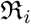 from the positive *x*_3_ axis. In this work it is assumed that 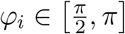 for each segment in the root system. This means that in proving (B.1) for *H_i_*(*c_a_*; *x*), it is only necessary to consider case 1: 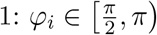 and case 2: 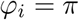.

**Case 1,** 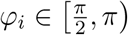: Completing the steps detailed in (A.1) yields the rotation matrix,

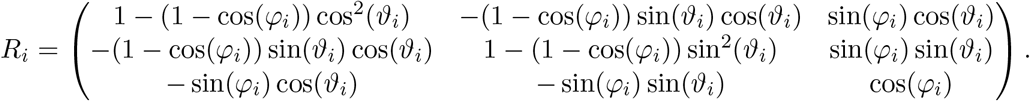

Using 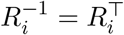, the matrix 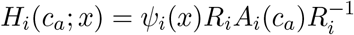 can be computed. For all 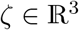 it is then the case that

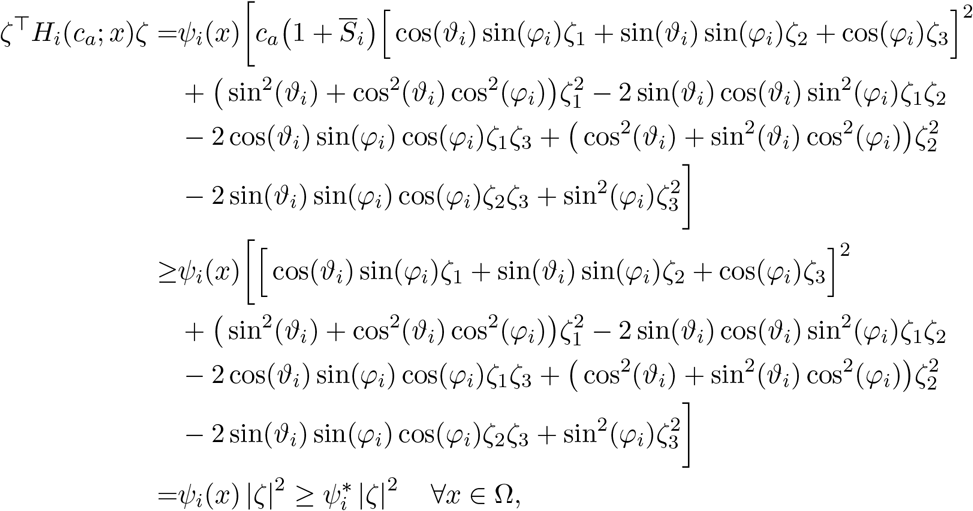

where 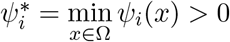.

**Case 2,** 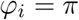: In this scenario the heterogeneity matrix associated to the segment is

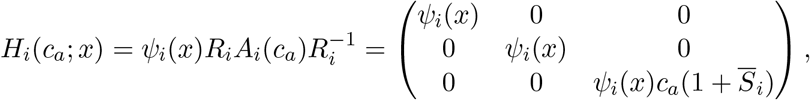

and as a result 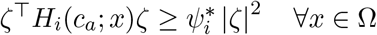 and 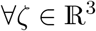. For any root segment 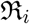 the matrix *H_i_* is therefore uniformly positive-definite. Since the sum of uniformly positive-definite matrices is also uniformly positive-definite, there must then exist λ > 0 such that *ζ*^⊤^*H*(*c_a_*; *x*)ζ ≥ λ |*ζ*|^2^ for all *x* ∈ Ω and 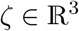.

## C Lipschitz continuous nondecreasing retention function (A1)

From (2.3) it can be seen that

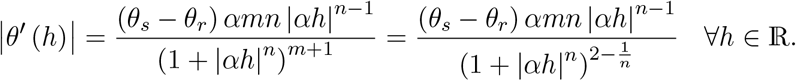

With the exact values stated in Section 3 for *θ_r_*, *θ_s_*, *α* and *n*, it can be seen that 0 < (*θ_s_* — *θ_r_*)*αmn* < 1 and 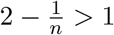. It then follows that

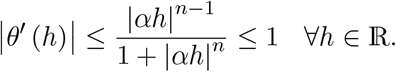

In order to show that the retention function is nondecreasing, the domain of *θ* is first restricted to the physical situation upon which this paper focuses, namely an unsaturated soil. This translates as nonpositive pressure head *h* ≤ 0 and it can then be seen from (2.3) that

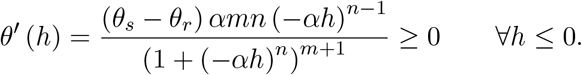

## D Condition (A2) for the regularised *κ_ε_*

The function *κ* given in (2.5), is bounded above by 1 but it is not Lipschitz continuous in *h* nor is it bounded below by a value greater than 0. In order to rectify this, a regularisation of *κ* was constructed and denoted by *κ_ε_*, where 0 < *ε* < (*θ_s_* — *θ_r_*). The first step in this regularisation is to define *θ_ε_* as follows:

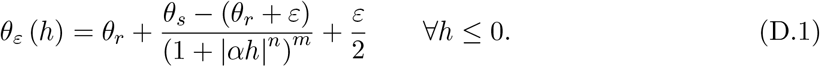

It can be seen that *θ_ε_* is non-decreasing for *h* ≤ 0 and retains the Lipschitz continuity of *θ*. However, unlike *θ*, *θ_ε_* exhibits the following limiting behaviour in *h* ≤ 0:

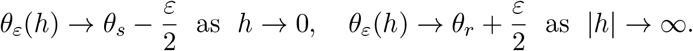

Then *κ_ε_* is defined as

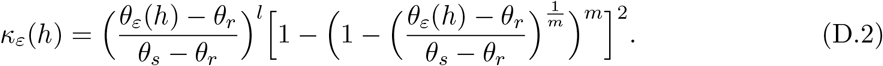

To show Lipschitz continuity of *κ_ε_* it can first be seen that 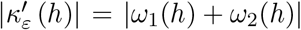 for *h* ∈ (-∞, 0], where:

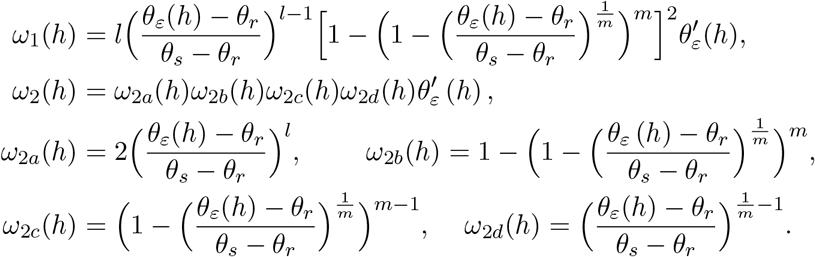

Direct calculations imply that for all *h* ∈ (-∞, 0]:

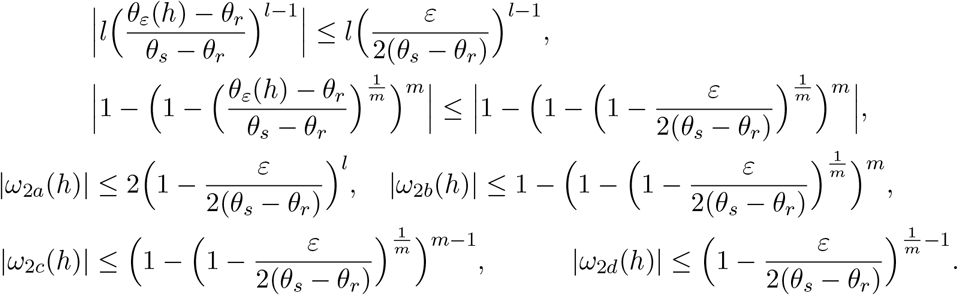

Combining the results above with the boundedness of 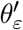 shows that 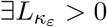 such that 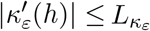 for all *h* ≤ 0, thus showing the Lipschitz continuity of *κ_ε_* for *h* ≤ 0.

Due to the limiting behaviour of *θ_ε_*, it is the case that for all *h* ∈ (-∞, 0],

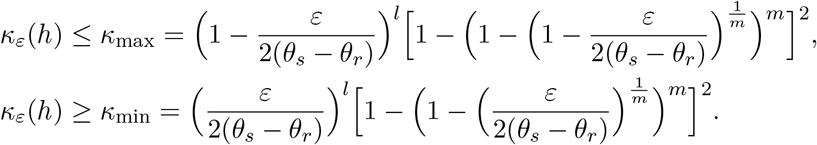

## References

Ahrens, J., B. Geveci, and C. Law (2005). Paraview: An end-user tool for large data visualization. The visualization handbook 717(8).

Alnæs, M. S., J. Blechta, J. Hake, A. Johansson, B. Kehlet, A. Logg, C. Richardson, J. Ring, M. E. Rognes, and G. N. Wells (2015). The fenics project version 1.5. Archive of Numerical Software 3(100).

Amenu, G. and P. Kumar (2008). A model for hydraulic redistribution incorporating coupled soil-root moisture transport. Hydrology and Earth System Sciences 12(1), 55–74.

Angers, D. A. and J. Caron (1998). Plant-induced changes in soil structure: processes and feedbacks. Biogeochemistry 42(1), 55–72.

Arbogast, T., J. Douglas, and U. Hornung (1990). Derivation of the double porosity model of single phase flow via homogenization theory. SIAM J Math Anal 21, 823–836.

Arbogast, T., M. Obeyesekere, and M. Wheeler (1993). Numerical methods for the simulation of flow in root-soil systems. SIAM J Numer Anal 30, 1677–1702.

Archer, N., J. N. Quinton, and T. Hess (2002). Below-ground relationships of soil texture, roots and hydraulic conductivity in two-phase mosaic vegetation in south-east spain. Journal of Arid Environments 52(4), 535–553.

Barczi, J.-F., H. Rey, S. Griffon, and C. Jourdan (2018). Digr: a generic model and its open source simulation software to mimic three-dimensional root-system architecture diversity. Annals of botany 121(5), 1089–1104.

Bastian, P., A. Chavarría-Krauser, C. Engwer, W. Jäger, S. Marnach, and M. Ptashnyk (2008). Modelling in vitro growth of dense root networks. Journal of Theoretical Biology 254(1), 99–109.

Beff, L., T. Günther, B. Vandoorne, V. Couvreur, and M. Javaux (2013). Three-dimensional monitoring of soil water content in a maize field using electrical resistivity tomography. Hydrology and Earth System Sciences 17(2), 595–609.

Bourgeat, A., S. Luckhaus, and A. Mikelić (1996). Convergence of the homogenization process for a dual-porosity model of immiscible two-phase flow. SIAM J Math Anal 27(6), 1520–1543.

Bourgeat, A., A. Mikelić, and A. Piatnitski (2003). On the double porosity model of a singlephase flow in random media. Asymptotic Analysis 34, 311–332.

Brent, R. P. (2013). Algorithms for minimization without derivatives. Courier Corporation.

Brochu, E., V. M. Cora, and N. De Freitas (2010). A tutorial on bayesian optimization of expensive cost functions, with application to active user modeling and hierarchical reinforcement learning. arXiv preprint arXiv:1012.2599.

Burkardt, J. (2014). The truncated normal distribution. Department of Scientific Computing Website, Florida State University, 1–35.

Carminati, A., A. B. Moradi, D. Vetterlein, P. Vontobel, E. Lehmann, U. Weller, H.-J. Vogel, and S. E. Oswald (2010). Dynamics of soil water content in the rhizosphere. Plant and soil 332(1), 163–176.

Danjon, F., D. Bert, C. Godin, and P. Trichet (1999). Structural root architecture of 5-year-old pinus pinaster measured by 3d digitising and analysed with amapmod. Plant and Soil 217(1), 49–63.

Darcy, H. P. G. (1856). Les Fontaines publiques de la ville de Dijon. Exposition et application des principes a suivre et des formules à employer dans les questions de distribution d’eau, etc.V. Dalamont.

Dewancker, I., M. McCourt, and S. Clark (2016). Bayesian optimization for machine learning: A practical guidebook. arXiv preprint arXiv:1612.04858.

Dorioz, J. M., M. Robert, and C. Chenu (1993). The role of roots, fungi and bacteria on clay particle organization. an experimental approach. In Soil Structure/Soil Biota Interrelationships, pp. 179–194. Elsevier.

Doussan, C., A. Pierret, E. Garrigues, and L. Pages (2006). Water uptake by plant roots: Ii-by plant roots: Ii–modelling of water transport in the soil root-system with explicit account of flow within the root system – comparison with experiments. Plant and Soil 283, 99–117.

Doussan, C., G. Vercambre, and L. Pagè (1998). Modelling of the hydraulic architecture of root systems: An integrated approach to water absorption—distribution of axial and radial conductances in maize. Annals of Botany 81(2), 225–232.

Dupuy, L., T. Fourcaud, A. Stokes, and F. Danjon (2005). A density-based approach for the modelling of root architecture: application to maritime pine (pinus pinaster ait.) root systems. Journal of Theoretical Biology 236(3), 323–334.

Dupuy, L., P. J. Gregory, and A. G. Bengough (2010). Root growth models: towards a new generation of continuous approaches. Journal of experimental botany 61(8), 2131–2143.

Frensch, J. and E. Steudle (1989). Axial and radial hydraulic resistance to roots of maize (zea mays l.). Plant Physiology 91(2), 719–726.

Gao, W., S. Schlüter, S. R. Blaser, J. Shen, and D. Vetterlein (2019). A shape-based method for automatic and rapid segmentation of roots in soil from x-ray computed tomography images: Rootine. Plant and Soil 441(1), 643–655.

Gerke, H. H. and M. T. Van Genuchten (1993). A dual-porosity model for simulating the preferential movement of water and solutes in structured porous media. Water resources research 29(2), 305–319.

Ghestem, M., R. C. Sidle, and A. Stokes (2011). The influence of plant root systems on subsurface flow: implications for slope stability. Bioscience 61(11), 869–879.

Gile, L., R. Gibbens, and J. Lenz (1995). Soils and sediments associated with remarkable, deeply-penetrating roots of crucifixion thorn (koeberlinia spinosazucc.). Journal of Arid Environments 31(2), 137–151.

Granier, A., N. Breda, P. Biron, and S. Villette (1999). A lumped water balance model to evaluate duration and intensity of drought constraints in forest stands. Ecological modelling 116(2-3), 269–283.

Harris, C. R., K. J. Millman, S. J. van der Walt, R. Gommers, P. Virtanen, D. Cournapeau, E. Wieser, J. Taylor, S. Berg, N. J. Smith, et al. (2020). Array programming with numpy. Nature 585(7825), 357–362.

Head, T., MechCoder, G. Louppe, I. Shcherbatyi, fcharras, Z. Vinícius, cmmalone, C. Schröder, nell215, N. Campos, T. Young, S. Cereda, T. Fan, rene rex, K. K. Shi, J. Schwabedal, carlosdanielcsantos, Hvass-Labs, M. Pak, SoManyUsernamesTaken, F. Callaway, L. Estève, L. Besson, M. Cherti, K. Pfannschmidt, F. Linzberger, C. Cauet, A. Gut, A. Mueller, and A. Fabisch (2018, March). scikit-optimize/scikit-optimize: v0.5.2.

Holzworth, D. P., N. I. Huth, P. G. deVoil, E. J. Zurcher, N. I. Herrmann, G. McLean, K. Chenu, E. J. van Oosterom, V. Snow, C. Murphy, et al. (2014). Apsim–evolution towards a new generation of agricultural systems simulation. Environmental Modelling & Software 62, 327–350.

Hornung, U. (1996). Homogenization and porous media. Springer.

Huang, L., P. Zhang, Y. Hu, and Y. Zhao (2015). Vegetation succession and soil infiltration characteristics under different aged refuse dumps at the heidaigou opencast coal mine. Global Ecology and Conservation 4, 255–263.

Johnson, M. S. and J. Lehmann (2006). Double-funneling of trees: Stemflow and root-induced preferential flow. Ecoscience 13(3), 324–333.

Jones, J. W., G. Hoogenboom, C. H. Porter, K. J. Boote, W. D. Batchelor, L. Hunt, P. W. Wilkens, U. Singh, A. J. Gijsman, and J. T. Ritchie (2003). The dssat cropping system model. European journal of agronomy 18(3-4), 235–265.

Leung, A., D. Boldrin, T. Liang, Z. Wu, V. Kamchoom, and A. Bengough (2018). Plant age effects on soil infiltration rate during early plant establishment. Géotechnique 68(7), 646–652.

Leung, A. K., A. Garg, and C. W. W. Ng (2015). Effects of plant roots on soil-water retention and induced suction in vegetated soil. Engineering Geology 193, 183–197.

Liang, T., A. Bengough, J. Knappett, D. MuirWood, K. W. Loades, P. D. Hallett, D. Boldrin, A. K. Leung, and G. Meijer (2017). Scaling of the reinforcement of soil slopes by living plants in a geotechnical centrifuge. Ecological Engineering 109, 207–227.

List, F. and F. A. Radu (2016). A study on iterative methods for solving richards’ equation. Computational Geosciences 20(2), 341–353.

Lourenço, S. D., K. Sassa, and H. Fukuoka (2006). Failure process and hydrologic response of a two layer physical model: implications for rainfall-induced landslides. Geomorphology 73(1-2), 115–130.

Mairhofer, S., S. Zappala, S. Tracy, C. Sturrock, M. J. Bennett, S. J. Mooney, and T. P. Pridmore (2013). Recovering complete plant root system architectures from soil via x-ray μ-computed tomography. Plant methods 9(1), 1–7.

Naseer, S., Y. Lee, C. Lapierre, R. Franke, C. Nawrath, and N. Geldner (2012). Casparian strip diffusion barrier in arabidopsis is made of a lignin polymer without suberin. Proceedings of the National Academy of Sciences 109(25), 10101–10106.

Noguchi, S., A. R. Nik, B. Kasran, M. Tani, T. Sammori, and K. Morisada (1997). Soil physical properties and preferential flow pathways in tropical rain forest, bukit tarek, peninsular malaysia. Journal of Forest Research 2(2), 115–120.

Noguchi, S., Y. Tsuboyama, R. C. Sidle, and I. Hosoda (1999). Morphological characteristics of macropores and the distribution of preferential flow pathways in a forested slope segment. Soil Science Society of America Journal 63(5), 1413–1423.

Passot, S., V. Couvreur, F. Meunier, X. Draye, M. Javaux, D. Leitner, L. Pages, A. Schnepf, J. Vanderborght, and G. Lobet (2019). Connecting the dots between computational tools to analyse soil-root water relations. Journal of experimental botany 70(9), 2345–2357.

Postma, J. A., C. Kuppe, M. R. Owen, N. Mellor, M. Griffiths, M. J. Bennett, J. P. Lynch, and M. Watt (2017). Opensimroot: widening the scope and application of root architectural models. New Phytologist 215(3), 1274–1286.

Radu, F. A., K. Kumar, J. M. Nordbotten, and I. S. Pop (2018). A robust, mass conservative scheme for two-phase flow in porous media including hölder continuous nonlinearities. IMA Journal of Numerical Analysis 38(2), 884–920.

Radu, F. A., I. S. Pop, and P. Knabner (2008). Error estimates for a mixed finite element discretization of some degenerate parabolic equations. Numerische Mathematik 109(2), 285–311.

Rasmussen, C. E. (2003). Gaussian processes in machine learning. In Summer school on machine learning, pp. 63–71. Springer.

Richards, L. A. (1931). Capillary conduction of liquids through porous mediums. Physics 1(5), 318–333.

Rodrigues, O. (1840). Des lois géométriques qui régissent les déplacements d’un système solide dans l’espace: et de la variation des cordonnées provenant de ces déplacements considérés indépendamment des causes qui peuvent les produire.

Schnepf, A., D. Leitner, M. Landl, G. Lobet, T. H. Mai, S. Morandage, C. Sheng, M. Zörner, J. Vanderborght, and H. Vereecken (2018). Crootbox: a structural-functional modelling framework for root systems. Annals of botany 121(5), 1033–1053.

Scholl, P., D. Leitner, G. Kammerer, W. Loiskandl, H.-P. Kaul, and G. Bodner (2014). Root induced changes of effective 1d hydraulic properties in a soil column. Plant and soil 381(1), 193–213.

Sidle, R. C., S. Noguchi, Y. Tsuboyama, and K. Laursen (2001). A conceptual model of preferential flow systems in forested hillslopes: Evidence of self-organization. Hydrological Processes 15(10), 1675–1692.

Simunek, J., N. Jarvis, M. van Genuchten, and A. Gärdenäs (2003). Review and comparison of models for describing non-equilibrium and preferential flow and transport in the vadose zone. J Hydrology 272, 14–35.

Simunek, J., M. T. van Genuchten, and M. Sejna (2005). The hydrus-1d software package for simulating the movement of water, heat, and multiple solutes in variably saturated media, version 3.0, hydrus software series 1. Department of Environmental Sciences, University of California Riverside, Riverside.

Slodicka, M. (2002). A robust and efficient linearization scheme for doubly nonlinear and degenerate parabolic problems arising in flow in porous media. SIAM Journal on Scientific Computing 23(5), 1593–1614.

Song, L., J. Li, T. Zhou, and D. Fredlund (2017). Experimental study on unsaturated hydraulic properties of vegetated soil. Ecological Engineering 103, 207–216.

Syme, W. (2001). Tuflow-two & onedimensional unsteady flow software for rivers, estuaries and coastal waters. In IEAust Water Panel Seminar and Workshop on 2d Flood Modelling, Sydney.

Teng, J., A. J. Jakeman, J. Vaze, B. F. Croke, D. Dutta, and S. Kim (2017). Flood inundation modelling: A review of methods, recent advances and uncertainty analysis. Environmental Modelling & Software 90, 201–216.

Thomee, V. (2006). Galerkin finite element methods for parabolic problems, Volume 25 of Springer Series in Computational Mathematics. Springer Berlin Heidelberg.

Valle, N., K. Potthast, S. Meyer, B. Michalzik, A. Hildebrandt, and T. Wutzler (2017). Modeling macropore seepage fluxes from soil water content time series by inversion of a dual permeability model. Hydrol. Earth Syst. Sci. Discuss., 1–31.

van der Heijden, G., A. Legout, B. Pollier, C. Bréchet, J. Ranger, and E. Dambrine (2013). Tracing and modeling preferential flow in a forest soil—potential impact on nutrient leaching. Geoderma 195, 12–22.

Van Genuchten, M. T. (1980). A closed-form equation for predicting the hydraulic conductivity of unsaturated soils 1. Soil science society of America journal 44(5), 892–898.

Virtanen, P., R. Gommers, T. E. Oliphant, M. Haberland, T. Reddy, D. Cournapeau, E. Burovski, P. Peterson, W. Weckesser, J. Bright, et al. (2020). Scipy 1.0: fundamental algorithms for scientific computing in python. Nature methods 17(3), 261–272.

Wilcox, B. P., D. D. Breshears, and H. Turin (2003). Hydraulic conductivity in a piñon-juniper woodland: Influence of vegetation. Soil Science Society of America Journal 67(4), 1243–1249.

Williams, D. (1991). Probability with martingales. Cambridge university press.

Wu, J., R. Zhang, and S. Gui (1999). Modeling soil water movement with water uptake by roots. PLant and Soil 215, 7–17.

Zhao, X., L. Xing, S. Shen, J. Liu, and D. Zhang (2020). Non-destructive 3d geometric modeling of maize root-stubble in-situ via x-ray computed tomography. International Journal of Agricultural and Biological Engineering 13(3), 174–179.

